# Adaptive prediction intervals for polygenic risk scores reveal individual variation in genetic predictability

**DOI:** 10.1101/2025.10.13.682102

**Authors:** Chen Wang, Fan Wang, Malgorzata Bogdan, Marco Masala, Edoardo Fiorillo, Marcella Devoto, Francesco Cucca, Daniel W. Belsky, Iuliana Ionita-Laza

## Abstract

Polygenic risk scores (PRS) are widely used in post-GWAS analyses to predict complex traits across humans, animals, and plants, yet the uncertainty of these predictions is rarely quantified at the individual level. Here, we introduce a framework for individualized uncertainty quantification based on quantile regression and conformal prediction, enabling the construction of prediction intervals with guaranteed coverage under minimal assumptions. Quantile regression enables adaptive, individual-specific prediction intervals that capture asymmetry and allow interval widths to vary substantially across individuals based on genetic information alone. Applying this framework to 62 traits in the UK Biobank and the ProgeNIA/SardiNIA studies, we show that these intervals maintain valid coverage and reduce uncertainty in risk stratification compared to existing methods, driven by their adaptive construction. Prediction interval width correlates positively with age and BMI, indicating reduced genetic predictability in subsets of the population where genetic effects interact with environmental factors. Our results demonstrate that incorporating uncertainty is essential for interpreting polygenic predictions and provide a principled approach to distinguish individuals whose phenotypes are well explained by genetic predictors from those in whom non-genetic influences dominate.

## Introduction

PRS prediction is an important post-GWAS analysis whose goal is to predict an individual’s genetic risk for a particular trait or disease^1^. Unlike traditional risk factors, PRS can identify individuals at high or low risk from birth and therefore such individuals can, in principle, be prioritized or not for early lifestyle interventions. Despite substantial progress, PRS are not yet ready for broad clinical deployment for most traits^2^. One major issue is the limited prediction ability of such scores, which may be improved by incorporating additional information from biomarkers and risk factors for the disease of interest^3^. Another major limitation is that prediction uncertainty is not usually accounted for, which affects the performance of these scores in clinical risk stratification^4^. Uncertainty in PRS prediction is inherent due to uncertainty in SNP effect size estimates, possibly inaccurate modeling assumptions (e.g. linear additive model), unmodelled factors such as environment and gene-by-environment interactions, population structure, and chance. Therefore, the point estimate normally reported is insufficient and does not convey the underlying variability. Reporting and accounting for uncertainty is essential as that can affect medical decision making, while providing more transparency about the limitations of PRS for prediction purposes.

We investigate here the problem of uncertainty quantification for PRS from a novel perspective based on quantile regression (QR) that is a leading methodology for constructing individualized prediction intervals and provides important insights beyond what can be learned from linear regression (LR) models^5–8^. In the usual PRS prediction task the aim is to predict the mean of a phenotype (*Y*) given the genotype profile of an individual (*X*) assuming a linear model. The resulting PRS is a linear combination of individual genotypes weighted by estimated genetic effects from a LR model in GWAS. This is essentially a point prediction, meant to reflect the typical or average phenotypic outcome given a particular genotype profile, i.e. *E*(*Y*|*X*). A natural alternative to LR for quantitative traits is based on QR that extends mean-based LR to the analysis of conditional quantiles of phenotype *Y* given genotypes *X*^7,8^, *Q*_*Y*_(*τ*|*X*) for any quantile level *τ* ∈ (0,1); when *τ* = 0.5, *Q*_*Y*_(*τ*|*X*) corresponds to the conditional median of *Y* given genotypes *X*. QR consists of fitting a series of LR models for any quantile level *τ* ∈ (0,1), providing estimates of quantile-specific regression coefficients. QR is an overall well-powered and flexible approach that allows detection of genetic variants with both additive and more complex non-additive effects^7–9^. While variants with additive effects are well detected by LR, those with more complex effects tend to show heterogeneity across quantile levels and are better detected by QR techniques^7,8,10^. Importantly, QR allows us to move away from point predictions to prediction intervals that naturally account for the uncertainty in the point prediction (**Fig. 1(a)**). We further combine the QR methodology with conformal prediction^11^, a machine learning technique to build prediction intervals that helps improve coverage of the regular QR prediction intervals.

**Figure 1:**
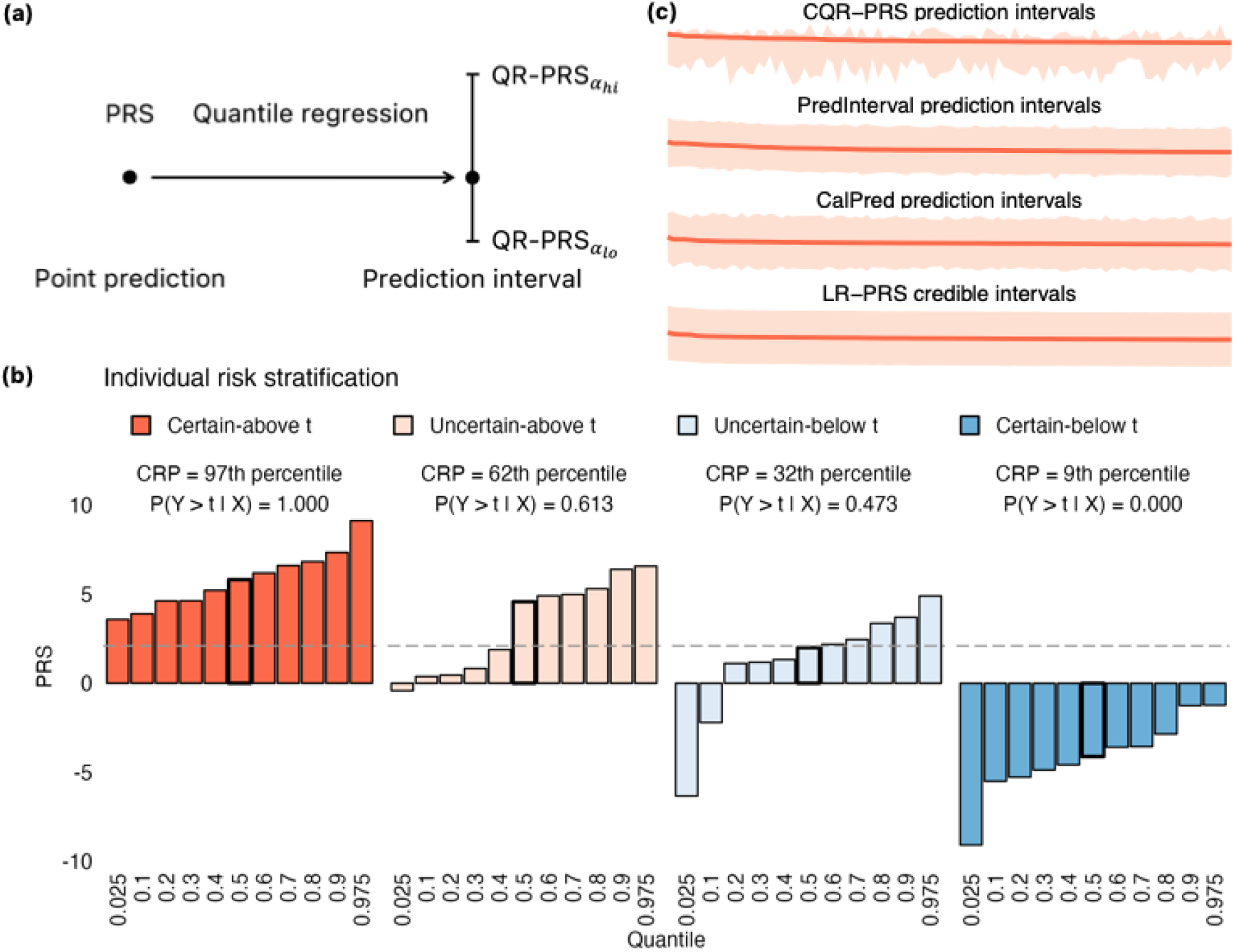
From point prediction to interval prediction for PRS. **(a)** Quantile regression naturally moves the emphasis from point prediction in conventional PRS to prediction interval that gives predictions of lower and upper quantiles of the conditional phenotype distribution. **(b)** Four individuals, each in a different risk category based on their predicted C-reactive protein (CRP) quantiles across a grid of quantile levels *τ* ∈ (0.025,0.975) (each bar corresponds to a predicted quantile). The threshold, *t* = 90^th^ percentile of QR-PRS_0.5_(top 10%), is shown as a gray horizontal line. For every individual we also report their actual CRP percentile and *P*(*Y* > *t*|*X*). **(c)** Prediction intervals (CQR-PRS_0.025_, CQR-PRS_0.975_), 95% prediction intervals from PredInterval and CalPred, and 95% credible intervals from LR-PRS are shown for top 100 individuals in uncertain-above categories for CQR-PRS, PredInterval, CalPred, and LR-PRS respectively.

Initial approaches to uncertainty quantification for PRS primarily concentrated on estimating uncertainty around the predicted phenotypic mean due to sampling uncertainty in the GWAS effect size estimates used to construct the polygenic score^4^. Subsequent approaches have extended uncertainty quantification to capturing variability throughout the phenotype distribution^12,13^. A key advantage of our approach that is primarily due to the use of QR is that it enables prediction intervals that vary in length across individuals and that are asymmetric based on genotype data alone. Such individualized prediction intervals can lead to more confident risk stratification by allowing some individuals’ entire intervals to lie above or below a high-risk threshold, something that is more difficult to achieve with less varying, symmetric LR-based intervals. Although recent methods such as CalPred^13^ can produce intervals of varying lengths across individuals (while remaining symmetric), they achieve this by explicitly incorporating individual-level non-genetic covariates. Individualized prediction intervals also enable differentiation between individuals with narrow intervals, whose trait levels may be largely captured by genetic predictors, and those with wider intervals, where non-genetic or environmental influences are likely more substantial. We refer to this new framework as CQR-PRS, for Conformalized Quantile Regression-PRS.

We illustrate CQR-PRS in simulations and applications to 62 quantitative traits using the UK Biobank (UKBB) data^14^. We include a detailed analysis for C-reactive protein, a moderately heritable inflammatory biomarker, with substantial contributions from both genetics and environmental factors. We validate the results in applications to an independent study, ProgeNIA/SardiNIA, a longitudinal study of a cohort of Sardinian subjects with many continuous phenotypes overlapping the traits analyzed in UKBB. In both simulations and real data applications, we compare performance with existing methods for interval construction.

## Methods

### CQR-PRS and prediction interval construction

At the core of CQR-PRS lies quantile regression, followed by conformal prediction to construct prediction intervals. CQR-PRS builds on the pruning + thresholding (P+T) method^15,16^, a popular PRS construction in the LR-based PRS literature, but employs QR instead of LR methodology. We acknowledge that this is a simple approach and more advanced genome-wide regression methods based on joint fitting of genetic variants genome-wide are preferable; however, such approaches are currently under development in the statistics literature and a biobank-scale application would require considerable additional engineering efforts and computational resources, which is currently challenging in practice^17^. In what follows we assume we have separate training/validation/test datasets, with details on these datasets being provided during specific applications.

#### 1) Pruning

Pruning is performed to reduce correlation among variants based on linkage disequilibrium (LD) estimated from the training data with a threshold of *r*^2^ < 0.1.

#### 2) Thresholding

A significance threshold is applied to filter out variants with potentially no effect on the trait. A simple threshold of 0.05 is applied to the quantile-specific QR p-values^7^ (see below on how to obtain these p-values using tools from QR). Note that this thresholding step can select different sets of variants at different quantile levels, as some variants with heterogeneous effects will only be significant at certain quantiles.

#### 3) QR-PRS computation

QR-PRS for each individual in the test dataset is calculated as the sum of the weighted effect alleles across *M* SNPs selected by pruning and thresholding above. The quantile-specific effect sizes estimated from the training data are used as weights of the selected variants, reflecting their contribution to the trait. For individual *i* and *M* SNPs with indices *j* = 1, …, *M* selected at quantile level *τ* we compute 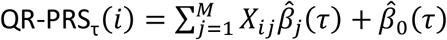, where 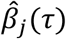 is the estimated coefficient for the *j*th variant at quantile level *τ, X*_*ij*_ is the corresponding genotype and 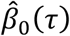 is the quantile-specific intercept. Note that we want to ensure quantile non-crossing, i.e. that the predicted quantiles QR-PRS_τ_(*i*) at quantile levels *τ* < *τ*^′^ are monotonic. We use here a common post-hoc adjustment based on sorting the predicted quantiles at a fine set of quantiles that ensures monotonicity^18,19^. A subtle point is that when variant effects *β*(*τ*) differ across quantiles *τ*, the QR-based intervals implicitly reflect underlying interactions including gene-by-environment (or gene-by-gene interactions). This adaptivity makes QR-PRS intervals biologically meaningful, as they encode heterogeneity in genetic effects rather than treating all genotypes as having constant effects across the trait distribution.

To estimate quantile-specific coefficients *β*(*τ*) used in steps 2 and 3 above, we fit linear QR models on the training data. Specifically, for the *j*th variant and specific quantile level *τ* ∈ (0,1) we fit the following conditional QR model:

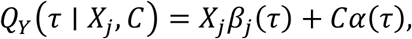

where *C* corresponds to covariates we want to adjust for; *β*_*j*_ (*τ*) and *α*(*τ*) are quantile-specific coefficients and can be estimated by minimizing the pinball loss function^20^. This optimization problem can be solved efficiently using existing tools. For testing, we employ a commonly used hypothesis testing tool in QR, the rank score test^21,22^ to test the null hypothesis *H*_0_: *β*_*j*_ (*τ*) = 0 and obtain p-values for individual SNPs at any given quantile level *τ*.

By fitting these models, we will be able to predict low and high phenotypic quantiles and build plug-in prediction intervals for each individual *i* of the form 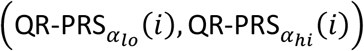. One potential issue is that plug-in QR prediction intervals are not guaranteed correct coverage due to various factors such as model misspecification, imprecision in estimation of conditional quantiles, and unmodelled important factors such as environment and gene-by-environment interactions. We discuss next conformalized versions of the plug-in intervals.

#### Conformal prediction intervals

Conformal prediction is a machine learning technique used to construct prediction intervals with valid coverage under minimal assumptions. By calibrating model predictions using held-out data, conformal methods can account for unmodelled sources of variability in the phenotype. As a result, conformal prediction has been widely applied across machine learning tasks to provide reliable uncertainty quantification^11^. A variant of conformal prediction has also been used previously to provide uncertainty for binary classification based on PRS prediction^23^. Combining conformal prediction with QR is a promising approach as it can lead to more realistic prediction intervals that are adaptive to heteroskedasticity in the data (due to their use of QR), while providing finite sample coverage guarantees^5^. We therefore build conformal versions of the plug-in prediction intervals above using split conformal prediction and refer to them as (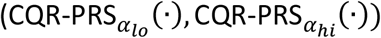).

Specifically, we fit QR models on the training data for some upper and lower quantile values *α*_*lo*_ and *α*_*hi*_. We then compute conformity scores on a separate calibration set 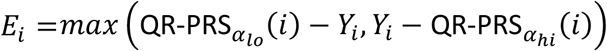 for each individual *i* in the calibration set. Given a new individual *n* + 1 we construct the conformalized prediction interval for *Y*_*n*+1_ as 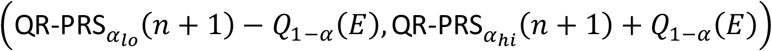, where 1 − *α* is the desired coverage level and *Q*_1 −*α*_(*E*) is the (1 − *α*)*th* empirical quantile of conformity scores *E*. This construction leads to prediction intervals with marginal coverage guarantee^11^. Note that the conformalization step extends the plug-in intervals 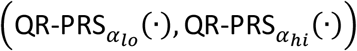 by the same amount *Q*_1−*α*_(*E*) in both directions. Consequently, the original QR-based intervals largely determine key properties of the conformalized intervals, including their asymmetry and the variation in length across individuals as discussed in this paper. The conformalization step serves only to guarantee correct coverage.

### Computation of *P*(***Y*** > ***t***|***X***)

Given the ability to estimate conditional quantiles *Q*_*Y*_(*τ*|*X*) at different quantile levels *τ* as described above, we can then compute *P*(*Y* > *t*|*X*) = 1 − *F*(*t*|*X*) for any threshold of interest *t*, where *F*(*Y*|*X*) is the cumulative distribution function (CDF). Specifically, the conditional quantile function is related to the CDF by the following formula: *Q*_*Y*_(*τ*|*X*) = inf{*y*: *F*(*y*|*X*) ≥ *τ*}, or equivalently 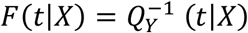. Therefore, given *Q*_*Y*_(*τ*|*X*) for a grid of quantile levels *τ* ∈ (0,1), we can use a smooth interpolation method such as b-splines^24^ to approximate the CDF.

We compare our QR-based framework with three methods in the literature based on LR, as described next.

### LR-PRS and credible interval construction

Uncertainty quantification for PRS based on LR (LR-PRS) is computed following the approach described in Ding et al.^4^ based on LDpred2^25^, a Bayesian method to estimate the posterior effect size distribution of each SNP by accounting for LD. Specifically, we first ran LDpred2 per autosomal chromosome with a grid of different values for several parameters, including heritability (0.7, 1.0, and 1.4 times the heritability estimated with LD score regression), polygenicity (evenly spaced on a logarithmic scale between 10^-4^ and 1), and sparsity (1 or 0 indicating whether or not a sparse model is applied) using a training dataset, as in Ding et al.^4^. The model (parameter set) with the highest R^2^ between the covariate-adjusted phenotype and the genetic prediction in a separate validation dataset was selected as the best PRS model to estimate the LR-PRS. For the optimal parameter setting, we ran LDpred2 a second time on the training dataset, with the Markov Chain Monte Carlo (MCMC) sampler with 100 burn-in iterations to produce B=500 posterior samples of effect sizes. Then, for each individual in the test dataset we computed a PRS estimate from each posterior sample to approximate the posterior distribution of PRS for that individual and empirically computed a *ρ*-level credible interval. defined by the 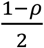 and 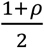 quantiles of the posterior distribution. We transformed the uncertainty estimates of the polygenic scores into corresponding phenotype uncertainty by adding the residual error variance, given by 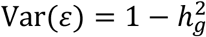, where 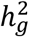 denotes the SNP-based heritability.

### CalPred and prediction interval construction

Prediction accuracy and uncertainty of PRS are affected by various contexts. CalPred is a calibration-based framework for constructing PRS prediction intervals that accounts for heterogeneity in prediction performance across individual contexts. CalPred models the residual phenotypic variance as a function of individual-level contexts, allowing prediction uncertainty to vary across individuals. We used the LDpred2-based LR-PRS as the point prediction and applied CalPred to model the residual variance using contextual variables including age, sex, batch, and the first 10 principal components in the calibration dataset. The fitted CalPred model was then applied to the test set to estimate the context-dependent prediction variance 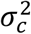. Prediction intervals for the test set were constructed using (LR-PRS − *z*_*ρ*_*σ*_*c*_, LR-PRS + *z*_*ρ*_*σ*_*c*_), where *z*_*ρ*_ is the standard normal quantile for the desired *ρ*-level prediction interval (for example, *z*_*ρ*_ = 1.96 for *ρ* = 95%).

### PredInterval and prediction interval construction

PredInterval is a non-parametric method based on the CV+ framework^26^ designed to construct calibrated prediction intervals for PRS-based phenotype prediction^12^. Like our proposed method, PredInterval also belongs to the class of conformal prediction approaches. To make results comparable with CQR-PRS, we built the PRS in PredInterval based on the P+T construction and LR model, using the same settings in terms of pruning and thresholding as in CQR-PRS (we have additionally considered using the PRS construction based on LDpred2 in LR-PRS above, but results remain qualitatively similar). Five-fold cross-validation (CV) was used to compute out-of-fold prediction residuals from the training data. The PredInterval training set is equivalent to the combined training and validation/calibration sets used by LR-PRS, CQR-PRS, and CalPred. Using the CV+ procedure implemented in PredInterval, we combined the model predictions (from the PRS) and the corresponding residuals to construct *ρ*-level prediction intervals for the independent test dataset.

### Interval asymmetry

We refer to an interval as being asymmetric if it is unbalanced around the point prediction (i.e. predicted median for CQR-PRS and predicted mean for LR-based methods). We quantify the extent of asymmetry by the ratio comparing how far the lower bound of a prediction interval is from the point prediction versus how far the upper bound is. If the interval extends equally above and below the prediction, the ratio equals one, meaning the interval is symmetric. When the ratio is less than one, the upper part of the interval is wider, indicating greater uncertainty toward higher predicted values. When the ratio is greater than one, the lower part of the interval is wider, showing more uncertainty toward lower predicted values. Allowing asymmetric prediction intervals is valuable in practice because it enables the interval to adapt to skewed or heteroskedastic data, providing more accurate and efficient coverage while reflecting the true uncertainty of the outcome.

### Simulation settings

We utilized the unimputed genotype data from UKBB white British samples. For quality control, we excluded individuals with more than 10% missing genotypes and retained SNPs with a minor allele frequency (MAF) greater than 5%, a genotyping rate of at least 90%, and a Hardy-Weinberg equilibrium p-value greater than 10^-6^. After quality control, 326,377 unrelated individuals and 340,091 SNPs were retained for downstream simulations. The dataset was partitioned into three disjoint sets: a train set comprising 240,000 individuals, a validation (for LDpred2-based LR-PRS) or calibration dataset (for CQR-PRS) comprising 60,000 individuals, and a test set with 26,377 individuals. PRS and prediction intervals were modeled using the training and validation/calibration sets. PredInterval was implemented using five-fold CV. We combined the training and calibration sets to form the training data (N = 300,000 individuals) for PredInterval. Therefore, each PredInterval fold used 240,000 individuals for training and 60,000 individuals for calibration, matching the sample sizes used in LR-PRS and CQR-PRS. Model performance including coverage, interval length, and interval asymmetry was evaluated in the independent test set for all methods.

In a first step, LR and QR GWAS were performed using the training set. LR-PRS was constructed using LDpred2 trained on the training set and optimized on the validation set. The optimized model was then retrained on the training set to generate posterior samples of SNP effect sizes. These posterior samples were subsequently applied to the test set to compute PRS point estimates and *ρ*-level credible intervals. For CQR-PRS, a P+T PRS (LD pruning with r^2^<0.1 and significance threshold p < 0.05) was constructed on the training set using QR-GWAS summary statistics at each quantile level. In the calibration set, the same model was applied to compute conformity scores. The conformity scores were then used to derive the CQR prediction intervals for the test set.

We evaluate performance for the resulting intervals using simulations under two settings. We consider first a homogeneous setting where all causal variants are homogeneous, with same effect across quantiles. Then we consider a heterogeneous setting where a proportion of causal variants have heterogeneous effects across quantiles.

#### Homogenous settings

Given the genotype matrix **X**, genetic heritability 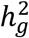, and proportion of causal variants *p*_causal_, standardized effects and phenotypes are generated as follows:

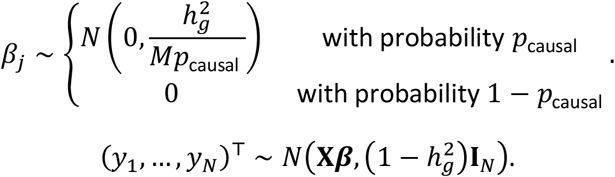

We simulate data under a range of parameters: 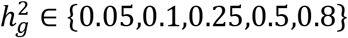 and *p* ∈ {0.001,0.01,0.1,1}, for a total of 20 simulation settings.

#### Heterogeneous settings

As above we generate standardized effect sizes and phenotypes as follows:

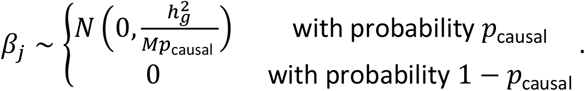

For *m* = *m*_1_ + *m*_2_ causal variants, we assume that *m*_1_ variants have homogenous effects and *m*_2_ variants have heterogeneous effects, i.e. local, non-zero effects only in the upper quantiles. Such local effects can reflect gene-by-environment interactions, where effects only manifest in certain environments that may correspond to upper quantiles, for example. For a variant *l* = (*m*_1_ + 1) … *m* with heterogeneous effects, *b*_*l*_(*τ*) is a local effect defined as

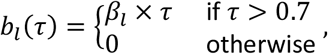

where *τ* ∈ (0,1) is a quantile level. Hence *b*_*l*_(*τ*) is non-zero only for quantile level *τ* > 0.7. We then generate the phenotype *Y* as follows:

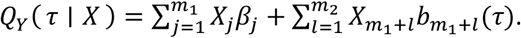

To generate *Y*, we use the inverse transform sampling. We simulate *β* using a range of parameters: and *p*_causal_ ∈ {0.001,0.01,0.1,1}. We assume *m*_1_ = *m*_2_ in our simulations.

## Results

## Simulations

Simulations below are based on the following: a train set comprising 240,000 individuals, a validation (for LR-PRS) or calibration dataset (for CQR-PRS) comprising 60,000 individuals, and a test set with 26,377 individuals. Each PredInterval CV fold uses 240,000 individuals for training and 60,000 individuals for calibration, matching the sample sizes used in LR-PRS and CQR-PRS. See more details in the “Simulation settings” section (Methods). Note that these simulations do not include any non-genetic covariate, so we do not evaluate CalPred in simulations.

We evaluate the empirical coverage of the constructed intervals under the two simulation scenarios described above for individuals in the independent test data. Specifically, we compute the proportion of individuals whose CQR-PRS prediction intervals (CQR-PRS_*α*_(·), CQR-PRS_1-*α*_(·)) for different values of *α* ∈ {0.025, 0.1, 0.2, 0.3, 0.4} overlap their true phenotype value. We use similar definitions for PredInterval and LR-PRS.

CQR-PRS is well calibrated across all settings (**Fig. 2, Fig. S1**). PredInterval exhibits slight overcoverage across scenarios, which is expected because the CV+ procedure is inherently conservative and produces intervals with guaranteed coverage regardless of model accuracy^5,26^. LR-PRS has overall close to correct coverage, with a tendency of undercoverage in sparse regimes (when the proportion of causal variants is low) and when the heritability is high.

**Figure 2:**
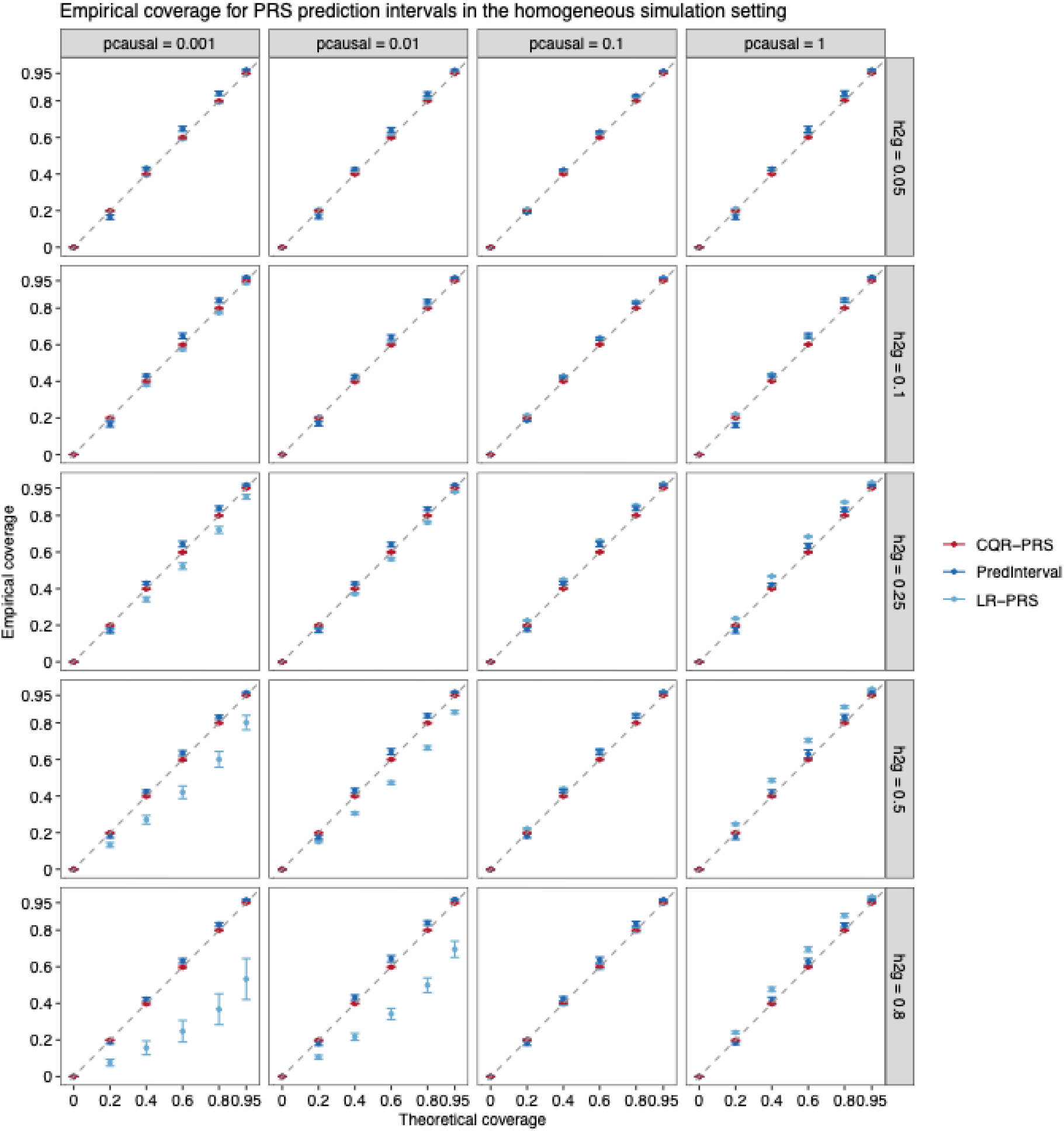
Empirical coverage vs. theoretical coverage for CQR-PRS, PredInterval and LR-PRS intervals in the homogeneous simulation setting. Results are shown for varying heritability and polygenicity parameters. Error bars represent the mean coverage ± standard deviation, calculated across 20 replicates.

We also compare interval lengths for the different methods in simulations (**Fig. S2-S3**). The lengths of the intervals tend to be smallest and with low variation for LR-PRS. PredInterval and CQR-PRS tend to have larger intervals with similar median size, with CQR-PRS showing larger variation in length especially for the heterogeneous settings. The larger variation in interval width for CQR-PRS is expected, as QR explicitly models conditional quantiles, allowing prediction intervals to adapt to local heteroscedasticity in the phenotype distribution.

Finally, we evaluated the asymmetry of intervals from each method. As expected, LR-PRS and PredInterval produce largely symmetric intervals, whereas CQR-PRS intervals exhibit substantial asymmetry, reflecting their reliance on QR and the separate estimation of lower and upper phenotypic quantiles (**Fig. S4-S5**).

### Applications to C-reactive protein in UKBB

We first show results on applications to C-reactive protein (CRP), an inflammatory biomarker with both genetic and environmental contributions. The training dataset for PRS derivation is the white British cohort (n=325,306 unrelated individuals and ∼8.6 million SNPs) in UKBB where the original LR and QR GWAS summary statistics are derived. The test dataset includes n=29,643 unrelated European samples in UKBB that are not used in the training or calibration steps. A calibration set comprising n=10,000 unrelated European ancestry individuals from UKBB was used to perform the conformal calibration for CQR-PRS and contextual calibration for CalPred. LR-PRS (LDpred2) utilized this dataset as a validation set to determine the optimized PRS model. PredInterval was implemented with five-fold CV using a dataset combining the aforementioned training and calibration datasets. The sample sizes of the training, calibration/validation, and test sets for the other traits analyzed in UKBB are summarized in **Table S1**.

We perform QR at different quantile levels *τ* ∈ (0.025, 0.975), adjusting for several covariates including age, sex, batch, and 10 principal components (PC) of genetic variation (**Fig. S6)**. The estimated quantile-specific coefficients are used to compute quantile specific PRS (QR-PRS_*τ*_) for every individual in the test data.

In addition to the typical LR-PRS distribution, we now obtain quantile-specific QR-PRS distributions across a grid of quantile levels *τ* ∈ (0.025,0.975) (**Fig. S7**; see **Fig. S19** for CRP in ProgeNIA/SardiNIA). QR-PRS_0.5_(corresponding to median regression) is highly correlated with the mean-based LR-PRS (**Fig. S8**; see **Fig. S20** for CRP in ProgeNIA/SardiNIA) as expected. CRP has similar positive but weak correlation levels with LR-PRS and QR-PRS for the central quantiles *τ* ∈ (0.4,0.6). While QR-PRS corresponding to the central quantiles are highly correlated among themselves (and with LR-PRS), correlations become weaker with QR-PRS at more extreme quantiles (e.g. *τ* = 0.025 or *τ* = 0.975).

#### Individualized prediction intervals

One key feature of CQR-PRS prediction intervals is that they are not constrained to be symmetric and are allowed to vary in length from individual to individual (**Fig. 1(c)**). Indeed, the CQR-PRS prediction intervals are asymmetric, especially for individuals with lower and higher point predictions, while intervals from the other methods are largely symmetric (**Fig. 3(a)-(b)**). Although other methods such as CalPred can produce intervals with varying length, it is due to explicit inclusion of non-genetic covariates. Notably, the asymmetry and uncertainty pattern is different for individuals with high vs. low predicted values. Specifically, for individuals with low predicted values, there is more uncertainty towards higher values, while for individuals with high predicted values, there is more uncertainty towards lower values. Note that for other traits such as height, which is highly polygenic and heritable with approximately normal distribution, the resulting prediction intervals exhibit lower levels of asymmetry (**Fig. S9**).

**Figure 3:**
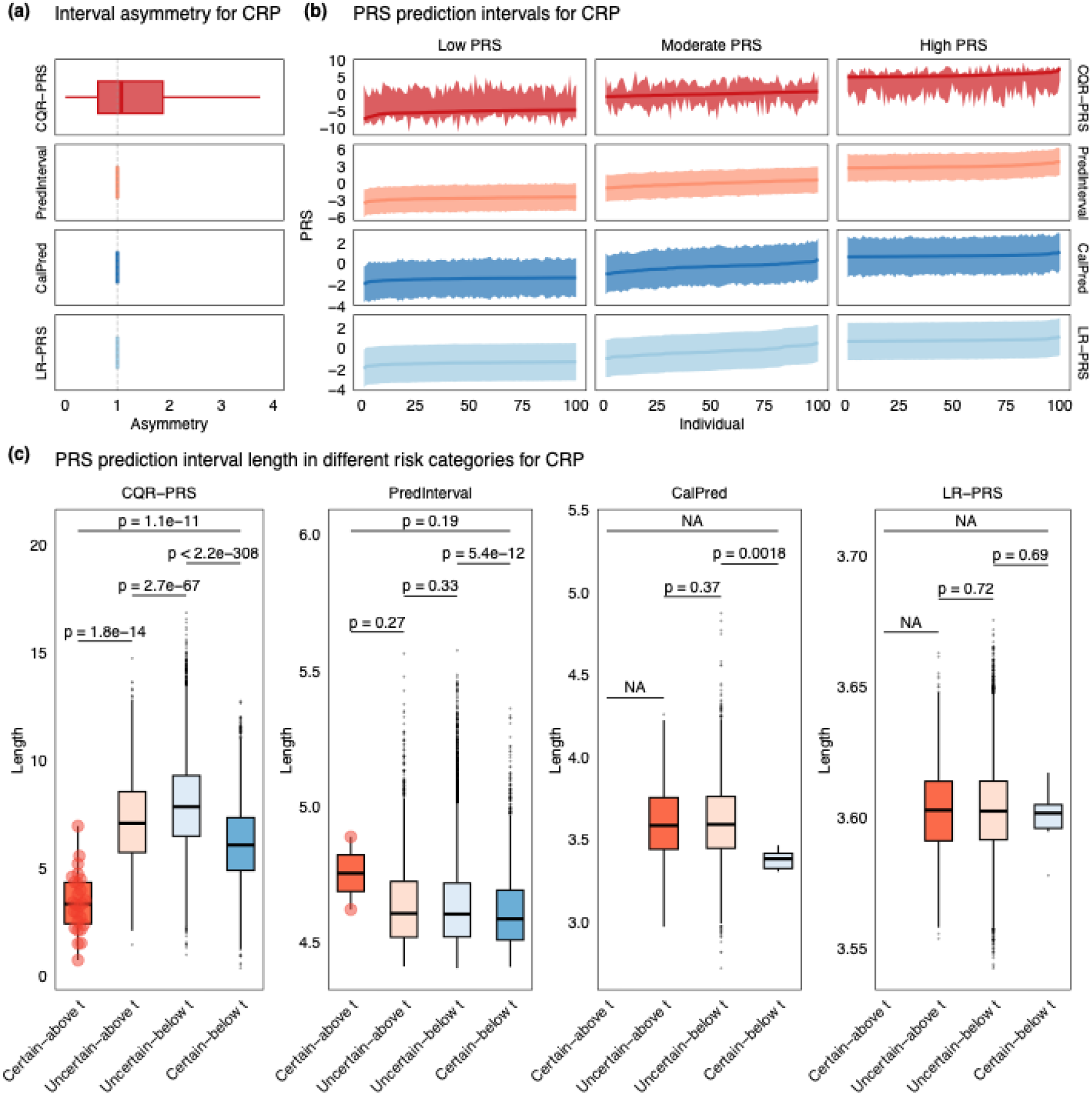
Individualized prediction intervals for CRP in UKBB. **(a)** CQR-PRS prediction intervals are asymmetric unlike those from PredInterval, CalPred, and LR-PRS. **(b)** Prediction intervals from CQR-PRS, PredInterval, CalPred, and LR-PRS for individuals with low, moderate, and high predicted values. Dots correspond to the QR-PRS_0.5_, PredInterval, CalPred, and LR-PRS point predictions. **(c)** Lengths of prediction intervals in different risk categories. Overlaid points on the ‘Certain-above *t*’ boxplots display individual data points, highlighting the notably small sample size in this category for the PredInterval method compared to CQR-PRS. The reported two-sided *p* values are from a Wilcoxon test comparing the length distribution in different categories.

Furthermore, individuals in the certain-above/certain-below *t* risk categories for CQR-PRS have narrower prediction intervals relative to individuals in the uncertain-above/uncertain-below *t* risk groups (**Fig. 3(c)**), and hence individuals with lower classification uncertainty have narrower prediction intervals. Furthermore, individuals with similar QR-PRS_0.5_values can have different prediction intervals. Such cases can arise if, for example, some of these individuals carry specific variants with effects only at upper or lower quantiles of the phenotype distribution (and negligible effects on the mean phenotype). Such heterogeneous effects across quantile levels may reflect underlying gene-by-environment or context-specific effects^7^. Therefore, the inclusion of such variants in the QR-PRS construction allows us to implicitly capture some unmodelled interactions or context-dependent associations. This is an important point because, while gene-by-environment interactions are assumed to play a crucial role in determining biomarkers and quantitative traits^27^, they are difficult to model explicitly given the large number of possible interacting factors and the fact that only few such factors can be assessed in any given study.

#### Risk classification

We use conformalized prediction intervals (CQR-PRS_*α*_(·), CQR-PRS_1-*α*_(·)) to perform more detailed risk stratification than possible based on just the single point prediction PRS. For example, if we consider a high-risk threshold *t* (e.g. the 90^th^ percentile of the QR-PRS_0.5_distribution), we can classify individuals into more refined risk categories, as proposed in Ding et al.^4^ (note that a similar classification is possible for the different methods discussed in this paper).

In particular, individual *i* with QR-PRS_0.5_above the high-risk threshold *t* can be classified as:

1. certain-above *t*: if (CQR-PRS_*α*_(*i*), CQR-PRS_1-*α*_(*i*)) is above *t*.
2. uncertain-above *t*: if (CQR-PRS_*α*_(*i*), CQR-PRS_1-*α*_(*i*)) contains *t*.

Similarly, individual *i* with QR-PRS_0.5_below the high-risk threshold *t* can be classified as:

1. uncertain-below *t*: if (CQR-PRS_*α*_(*i*), CQR-PRS_1-*α*_(*i*)) contains *t*.
2. certain-below *t*: if (CQR-PRS_*α*_(*i*), CQR-PRS_1-*α*_(*i*)) is below *t*.

We take *α* = 0.025 for illustration (unless otherwise specified). Examples of individuals in each of the four categories are shown in **Fig. 1(b)**. For each individual, we also compute tail probabilities *p* = *P*(*Y* > *t*|*X*) for a given threshold *t* (90^th^ percentile of QR-PRS_0.5_). For the four individuals in **Fig. 1(b)**, these probabilities correspond well to our intuition, i.e. certain-above *t*: *p* = 1; uncertain-above *t*: *p* = 0.61; uncertain-below *t*: *p* = 0.47; and certain-below *t*: *p* = 0.

Similar to Ding et al.^4^ we can calculate the proportion of individuals classified as “high-risk”, based on the point prediction QR-PRS_0.5_exceeding a threshold *t*, who fall into the certain-above *t* vs. uncertain-above *t* groups. Similarly, among individuals classified as “low-risk” (i.e. those with point prediction QR-PRS_0.5_< *t*), we compute the proportion belonging to the certain-below *t* vs. uncertain-below *t* groups. Overall, CQR-PRS yields the largest proportions in both the high-risk and low-risk groups (**Fig. 4(a)**) as we vary the nominal coverage level, although these proportions remain small when the target coverage is high (e.g. 95%). For example, among individuals with “high” QR-PRS_0.5_, i.e. at or above the 90^th^ (80^th^) percentile, only 0.9% (1.6%) are classified in the certain-above *t* group. CQR-PRS also identifies the largest number of individuals who are classified as certain-above *t* and with actual CRP values exceeding the 90^th^ (80^th^) percentile of the UKBB distribution (**Fig. 4(b)**). Similar trends hold for individuals in the low-risk group, although the proportions are much higher than for the high-risk category. For example, among those at or below the 90^th^ (80^th^) percentile of the QR-PRS_0.5_distribution, 30.8% (18%) are in the certain-below *t* category. This is important from a risk stratification perspective because it allows to identify individuals that may need less frequent screening or intervention compared to those at high risk.

**Figure 4:**
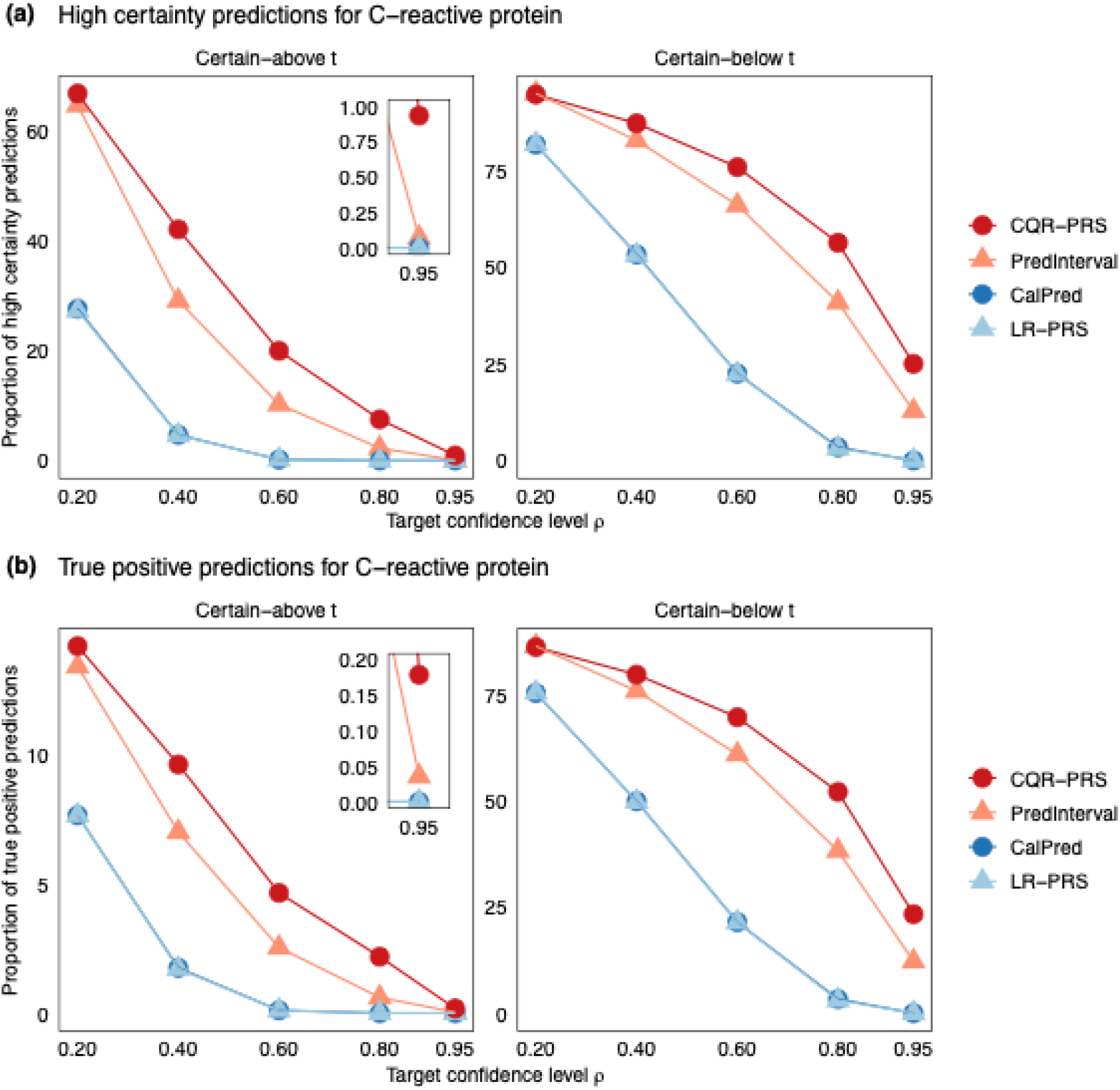
Proportion of high certainty and true positive predictions for CRP at different target coverage levels (*ρ* = 0.2, 0.4, 0.6, 0.8, 0.95) in UKBB. Proportion of high certainty predictions in panel **(a)** is calculated as # certain-above t / # above t and # certain-below t / # below t. Proportion of true positive predictions in panel **(b)** is calculated as # (certain-above t & trait above t) / # above t and # (certain-below t & trait below t) / # below t. The threshold *t* corresponds to the 90^th^ QR-PRS_0.5_/LR-based PRS or trait percentile depending on context.

Tail probabilities *P*(*Y* > *t*|*X*) have different distributions for individuals in different risk categories (**Fig. S10(a)**). The correlation between CRP and *P*(*Y* > *t*|*X*) is positive (**Fig. S10(b)**) but not particularly strong, likely due to environmental factors playing an important role for CRP. On the other hand, the correlation between QR-PRS_0.5_and *P*(*Y* > *t*|*X*) is stronger as expected (**Fig. S10(c)**). Additionally, the probabilities *P*(*Y* > *t*|*X*) from QR-PRS and LR-PRS are highly correlated (**Fig. S10(d)**).

### Applications to 62 traits

In addition to CRP, we applied QR-PRS to 61 other quantitative traits with a range of heritability values (**Table S1**). We show that across these diverse traits the conformal prediction intervals from CQR-PRS and PredInterval have valid coverage across all traits, while LR-PRS and CalPred intervals can suffer from undercoverage in certain cases (**Fig. 5**). These results are expected. Unlike conformal methods such as CQR-PRS and PredInterval that have theoretical coverage guarantees, there is no such guarantee for LR-PRS and CalPred.

**Figure 5:**
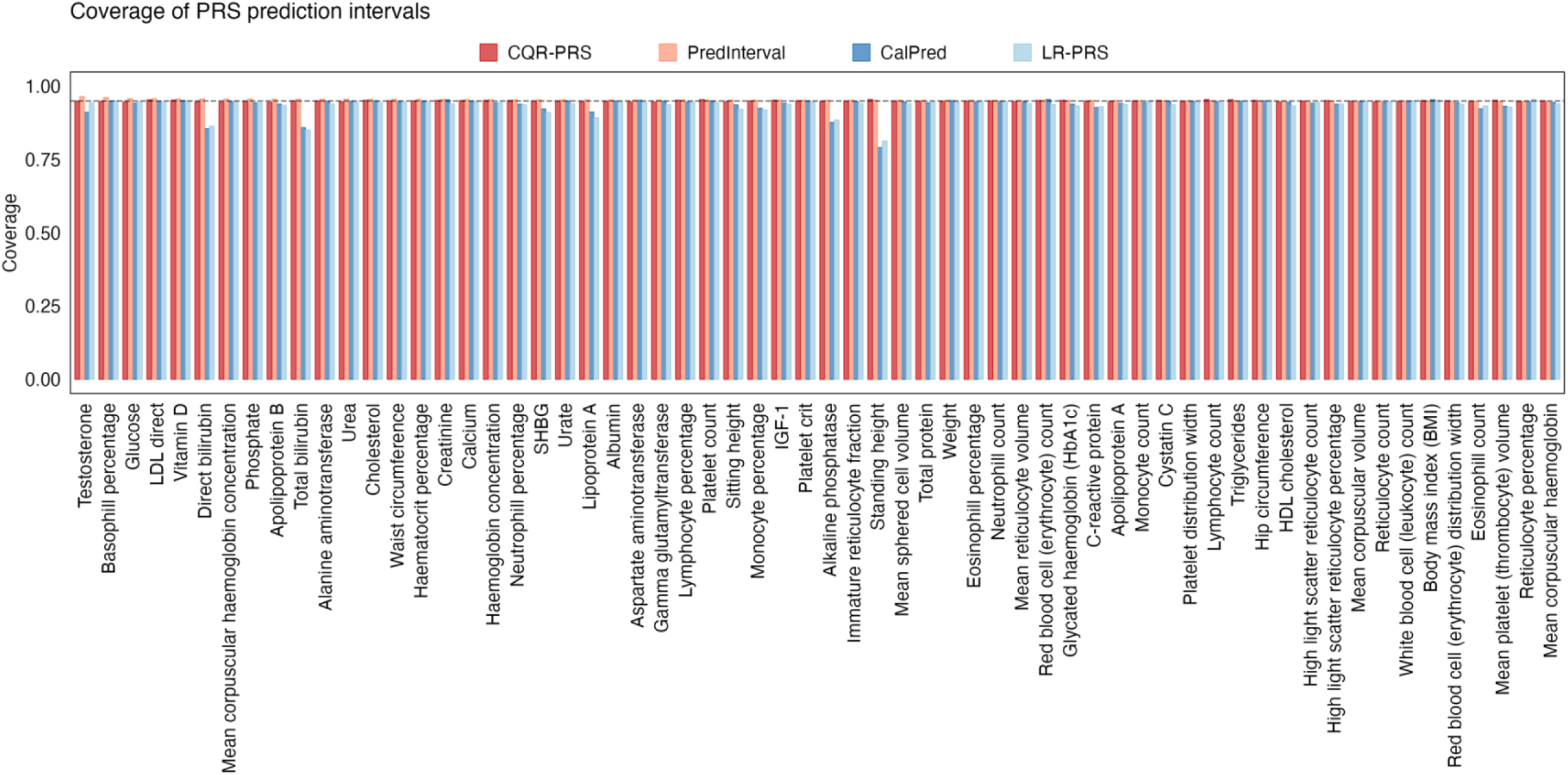
Coverage for prediction/credible intervals from CQR-PRS, PredInterval, CalPred and LR-PRS across 62 traits in UKBB. Coverage is computed as the proportion of individuals whose 95% level intervals overlap their true phenotype value.

Although, as expected, we observe a pattern of high uncertainty in risk stratification across all traits, CQR-PRS identifies a significantly greater number of individuals in the certain-above *t* and certain-below *t* categories compared to LR-PRS, CalPred and PredInterval (**Fig. 6(a), Fig. S11-S14**). Especially for the certain-below *t* category, the proportions of individuals identified by CQR-PRS are substantially higher than for LR-based methods across traits. This result is due to the adaptive nature of the CQR prediction intervals: they are asymmetric and vary in length much more than LR-based intervals (**Fig. S15-S16**). While CQR intervals might be wider on average, they dynamically adapt to the data. Compared to LDpred2 methods (such as CalPred and LR-PRS), CQR intervals are actually narrower relative to the overall variance of the PRS predictions. This relative precision allows CQR to produce a higher rate of definitive, actionable predictions for individuals falling strictly above or below the threshold (**Fig. S17**).

**Figure 6:**
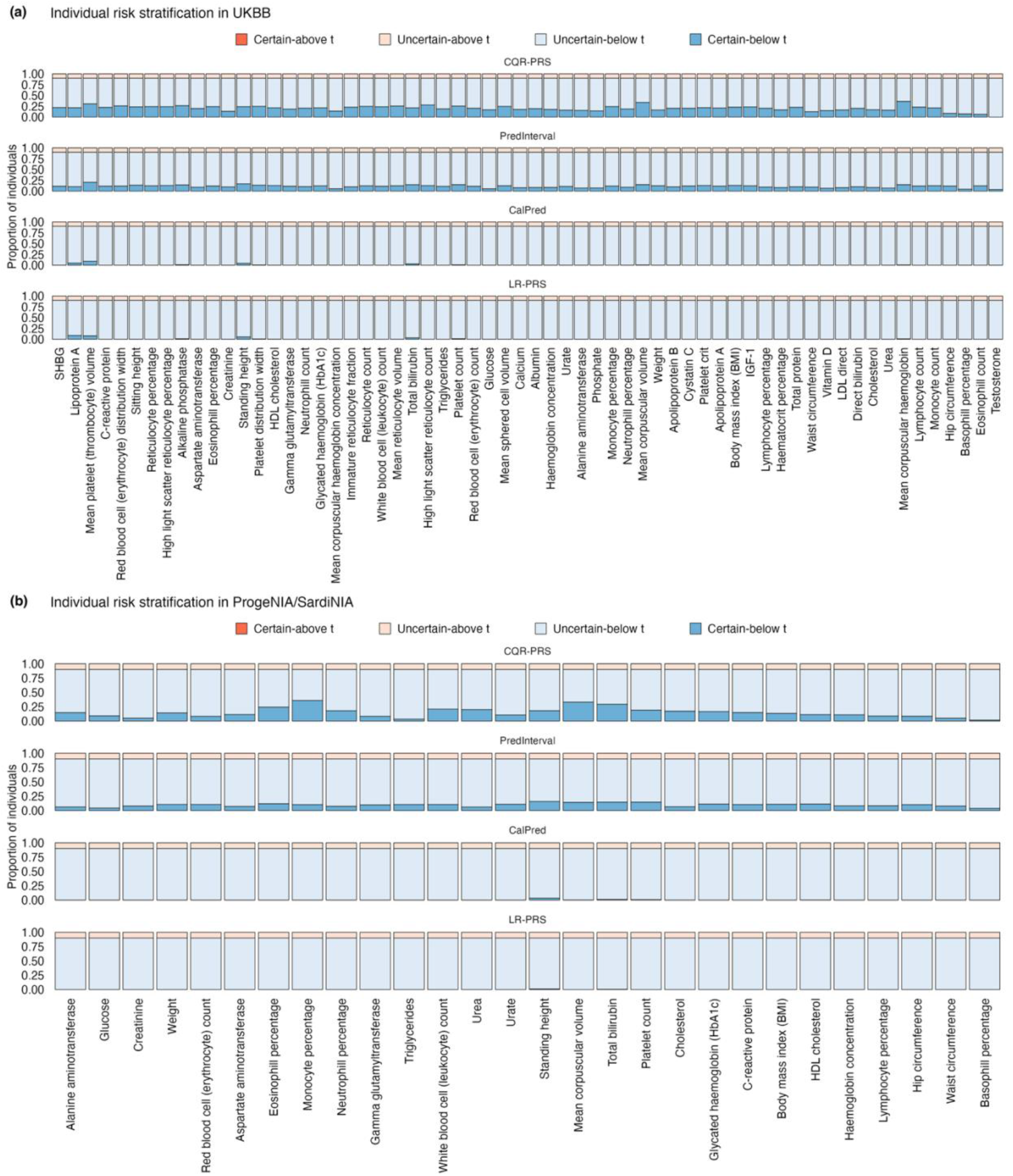
Risk stratification based on CQR-PRS, PredInterval, CalPred and LR-PRS across 62 traits. **(a)** Risk stratification across 62 traits in UKBB. **(b)** Risk stratification across 28 traits in ProgeNIA/SardiNIA. Stacked barplots correspond to proportion of individuals in the four risk categories certain-/uncertain-above and certain-/uncertain-below *t*. The threshold *t* corresponds to the 90^th^ QR-PRS_0.5_/LR-based PRS percentile.

Identifying individuals at low genetic risk is valuable in public health, as it helps ensure that expensive preventive strategies are directed toward those who will benefit most. Notably, there is significant and positive correlation between trait heritability and proportion of certain-below *t* predictions (for the low risk group), and proportion of true positive predictions for both high and low risk groups (**Fig. S18**).

#### Prediction interval width correlates with age

As noted above, a key advantage of QR-based prediction intervals is that their length naturally varies across individuals, even when only genetic information is included in their construction. For a given person, a narrow interval suggests that the model can predict that individual’s phenotype with relatively low uncertainty, indicating that their phenotype is more predictable from their genetic profile under the model. Conversely, a wider interval reflects higher individual-level uncertainty, implying a stronger influence of non-genetic or environmental factors, including possible gene-by-environment interactions. Thus, uncertainty quantification may highlight individuals whose phenotypes are more strongly influenced by environmental factors. Since environmental scores tend to accumulate over the life course, we examined the correlation between prediction interval width and age. We observed significant correlations (*p* < 0.05) between prediction interval width and age for 17/62 traits, with 16/17 of them being positive correlations (**Fig. 7(a)**). These positive correlations indicate that model uncertainty tends to increase with age, likely reflecting greater heterogeneity in non-genetic and environmental influences on trait with age. In contrast, fewer and less significant correlations were observed for LR-PRS and PredInterval, an expected result given the lower variation in length for LR-based methods described above and hence lower statistical power to identify significant correlations. CalPred exhibited strong correlations across traits, both positive and negative. However, unlike CQR, where age dependence arises implicitly from the data, CalPred explicitly parameterizes the error variance as a function of context (including age), making age-dependent variation in prediction intervals a built-in feature of the model.

**Figure 7:**
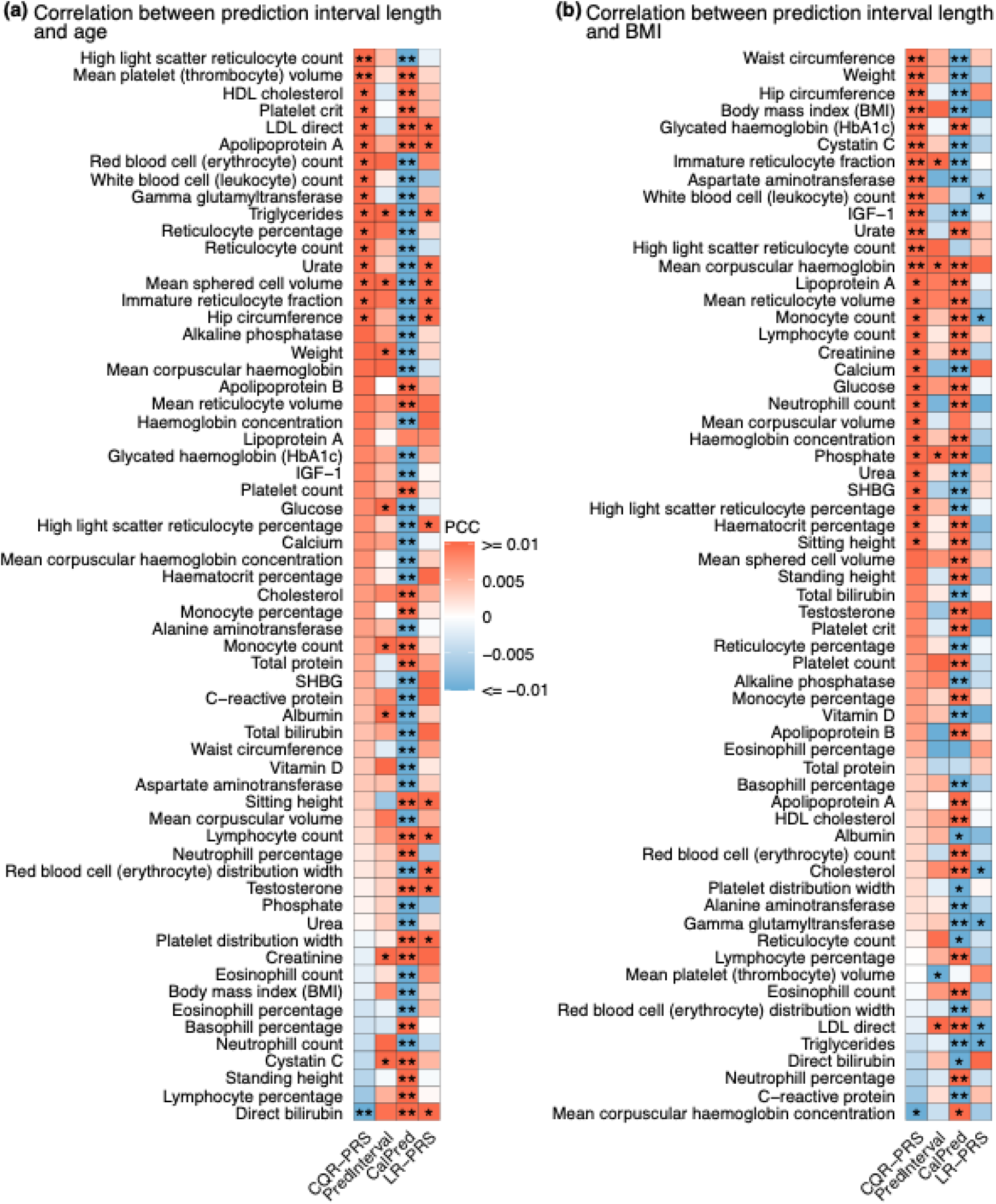
Correlations between interval length and (a) age; (b) BMI for 62 traits and four methods (CQR-PRS, PredInterval, CalPred and LR-PRS) in UKBB. Nominally significant correlations (*p* < 0.05) are indicated by “*”. Correlations significant after multiple testing correction (*p* < 0.05/(62 · 4)) are indicated by “**”.

#### Prediction interval width correlates with BMI

To further corroborate this result, we correlated prediction interval widths with several other factors including BMI, smoking status (current or ever), low physical activity and alcohol intake (current and ever drinker). Of these only BMI showed substantial correlations. Specifically, among the 62 traits, 30 exhibited significant correlations with BMI (*p* < 0.05), with 29/30 being positive correlations (**Fig. 7(b)**). Overall, these results suggest that wider intervals in QR-based prediction tend to occur in individuals with higher BMI. This pattern is consistent with the presence of gene-environment interactions or gene-environment correlations, under the assumption that BMI captures, at least in part, differences in environmental exposures or their variability. Notably, for adult height where environmental influences are relatively limited the prediction interval length shows no significant association with either age or BMI. Consistent with the age-related results above, PredInterval and LR-PRS show few significant correlations between interval length and BMI. In contrast, CalPred intervals exhibit strong correlations with BMI across most traits, in both positive and negative directions. Note that BMI is significantly correlated with age, sex, batch and multiple PCs used in the modeling of prediction intervals by CalPred.

Together with the similar associations between interval length and age observed above, these findings support the interpretation that increased environmental exposure or accumulated life-course variability is associated with reduced genetic predictability at older ages.

### Validations using the ProgeNIA/SardiNIA project data

We utilize a second, independent dataset from the ProgeNIA/SardiNIA project^28,29^, a longitudinal study of a cohort of Sardinian subjects (n=6,159) for validation. Of the 62 quantitative traits we analyzed in the UKBB, 28 are available to us from the ProgeNIA/SardiNIA project (**Table S1**). The results confirm the high uncertainty in risk stratification, and validate the finding that inference based on prediction intervals derived in the QR framework results in more individuals being assigned to the certain-above *t* and certain-below *t* risk categories relative to the counterpart LR frameworks (**Fig. 6(b), Fig. S21-S24**).

## Discussion

There is great interest in the human, plant and animal genetics communities to build trait prediction models based on genome-wide genotype data. For human genetics, the emphasis has been on employing PRS for risk stratification in clinical settings. However, this has proven to be challenging due to complexities underlying risk to disease including the important role of environmental effects and their complex interactions with genetic factors. Furthermore, although scientists are aware of the inherent imprecision of PRS estimates, this uncertainty is rarely reported or accounted for in analyses. However, taking uncertainty into account can lead to better risk stratification for individuals in high/low risk groups, while also providing more accurate information on the current limitations of PRS prediction.

We describe here a natural way to quantify individualized uncertainty based on building valid prediction intervals by combining QR and conformal prediction. The resulting intervals achieve correct marginal coverage under minimal assumptions and provide individualized assessments of uncertainty based on genotype data alone, as they can be asymmetric and vary in length across individuals. We further show that variation in interval length can reflect potential environmental influences at the individual level. From a clinical standpoint, wider prediction intervals can be informative, as they may indicate that unmodeled or modifiable environmental factors exert substantial influence on outcome variability within an individual. Such uncertainty highlights individuals whose true risk is more context-dependent and, therefore, potentially more responsive to lifestyle or environmental interventions. In this way, prediction interval width not only reflects the limits of purely genetic-based prediction but also helps identify patients for whom preventive strategies targeting modifiable exposures may yield the greatest benefit.

The computational cost of the proposed QR-based prediction interval construction is driven primarily by the QR GWAS step. For a typical dataset in our analyses comprising 325,306 individuals and 8,572,925 variants, this step required 216 CPU hours per quantile level, highlighting its computational intensity in large-scale settings. In contrast, downstream steps are comparatively efficient: clumping with PLINK2 required 3.7 CPU hours per quantile level, and computation of CQR-PRS point predictions and prediction intervals required only 0.2 CPU hours. These results indicate that the overall computational burden is dominated by the initial QR GWAS estimation, while subsequent steps are relatively lightweight. Importantly, there is scope for further optimization of the QR GWAS procedure, for example, through the use of smooth approximations to the pinball loss combined with gradient-based optimization methods, which we are currently investigating.

Although CQR-PRS is by design applicable to quantitative traits, it serves as a framework to illustrate how individualized uncertainty quantification can be achieved by constructing more realistic prediction intervals via QR. Moreover, biomarkers, drug response and intermediate cellular phenotypes^30^ are important in clinical medicine and pharmacogenomics. Similarly, for plants and animals constructing PRS for quantitative traits can be even more useful than for human studies for purposes such as selective breeding and conservation efforts^31–33^. Beyond quantitative traits, a future direction of interest will be to aggregate multiple QR-based PRS predictions at different quantile levels and for different disease-related endophenotypes or biomarkers to derive meta PRS scores for disease traits of interest with improved accuracy^34^, similar to ideas previously implemented in the context of LR^3,35^.

There are a few limitations of the proposed QR-PRS score and these will be the subject of future work. One inherent limitation is that QR requires individual level data to train the quantile regression models and thus cannot leverage existing summary statistics. However, given the large biobanks available across the world with individual level data and diverse biomarkers, including molecular, clinical and imaging biomarkers, and the increasing trend to collect quantitative traits across human, plant and animal studies, this is not a major limitation. Furthermore, summary statistics come with their own significant challenges and will likely lead to miscalibration in practice. Another limitation is that our current PRS construction is based on a simple pruning + thresholding method and has not been particularly optimized. A range of more sophisticated techniques including ridge regression and Bayesian regression methods^15,16^ could be incorporated in the quantile framework but that requires non-trivial effort in the context of quantile regression, especially from a computational perspective for biobank scale data^17^, and we leave that for future work. Such joint models may improve the efficiency of the resulting prediction intervals, potentially leading to shorter average interval lengths under conformal calibration. Finally, while conformal prediction guarantees marginal coverage (i.e. coverage averaged across all individuals), it does not guarantee conditional coverage. In particular, coverage within specific subgroups, such as individuals with high trait values, may deviate from the nominal level. We note that this limitation is not specific to our method but is inherent to other approaches considered in this paper such as PredInterval and LR-PRS. While CalPred can in principle achieve approximate conditional coverage, that will depend on the variance model being correct. Achieving conditional coverage is fundamentally hard, especially for conformal methods, and there is ongoing active work around constructing conditionally valid prediction intervals that could be useful in our settings in the future^36^. In addition to genetic factors, integrating non-genetic covariates such as age, sex, and other risk factors is expected to enhance the clinical utility of the resulting prediction intervals.

In summary, we describe a novel method to construct prediction intervals based on quantile regression and conformal prediction. By allowing asymmetric intervals of varying length, the proposed method can reveal where genetic predictions are more or less reliable and highlight individuals for whom environmental or non-genetic factors may play a larger role. Notably, the current model achieves varying length and asymmetry of prediction intervals using only genetic information. Overall, the proposed framework provides a new direction for uncertainty quantification and can serve as a foundational framework for future development of more complex models for constructing prediction intervals.

## Acknowledgements

This research has been conducted using the UK Biobank Resource under Application Number 41849. We thank Atlas Khan for help with the UKBB data processing. This research has been partially supported by NIH grant AG072272 and grant 2024-04735 from the Swedish Research Council.

## Data Availability

The individual-level genotype and phenotype data are available to approved researchers through the UK Biobank web portal at https://www.ukbiobank.ac.uk/. The LR and QR GWAS summary statistics for the biomarkers reported here are publicly available at https://doi.org/10.5281/zenodo.11095249 and https://doi.org/10.5281/zenodo.19503042.

## Code Availability

Scripts for CQR-PRS analyses are available at https://github.com/Iuliana-Ionita-Laza/QRPRS. The LR-PRS and PRS uncertainty estimation described in Ding et al. was performed using LDpred2 from https://github.com/privefl/bigsnpr and the code from https://github.com/bogdanlab/prs-uncertainty. CalPred analysis was performed using the code from https://github.com/KangchengHou/calpred. PredInterval was applied using the implementation available at https://github.com/xuchang0201/PredInterval. Scripts for QR GWAS analysis are available at https://github.com/Iuliana-Ionita-Laza/QRGWAS. The quantile regression was performed using the R package quantreg (https://cran.r-project.org/package=quantreg) version 5.94. The rank score test was performed using the R package QRank (https://cran.r-project.org/package=QRank) version 1.0. Analyses with these R packages were performed in R release 4.2.2. The linear regression analysis and P+T PRS were performed using the PLINK 2.0 (https://www.cog-genomics.org/plink/2.0/).

## Supplemental Material

**Figure S1:**
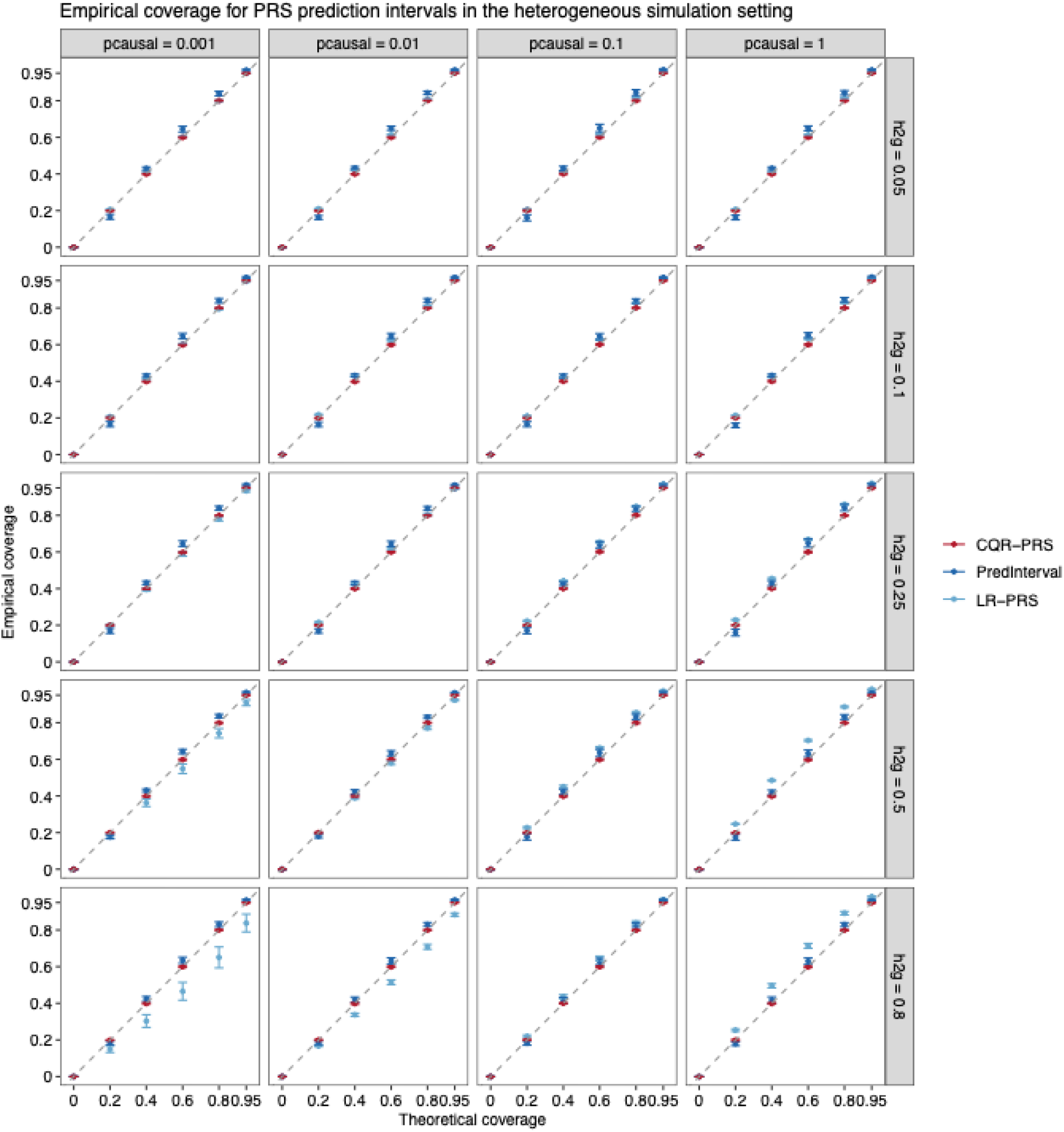
Empirical coverage vs. theoretical coverage for CQR-PRS, PredInterval, and LR-PRS intervals in the heterogeneous simulation setting (*m*_1_: *m*_2_ = 1: 1). Results are shown for varying heritability and polygenicity parameters. Error bars represent the mean coverage ± standard deviation, calculated across 20 replicates.

**Figure S2:**
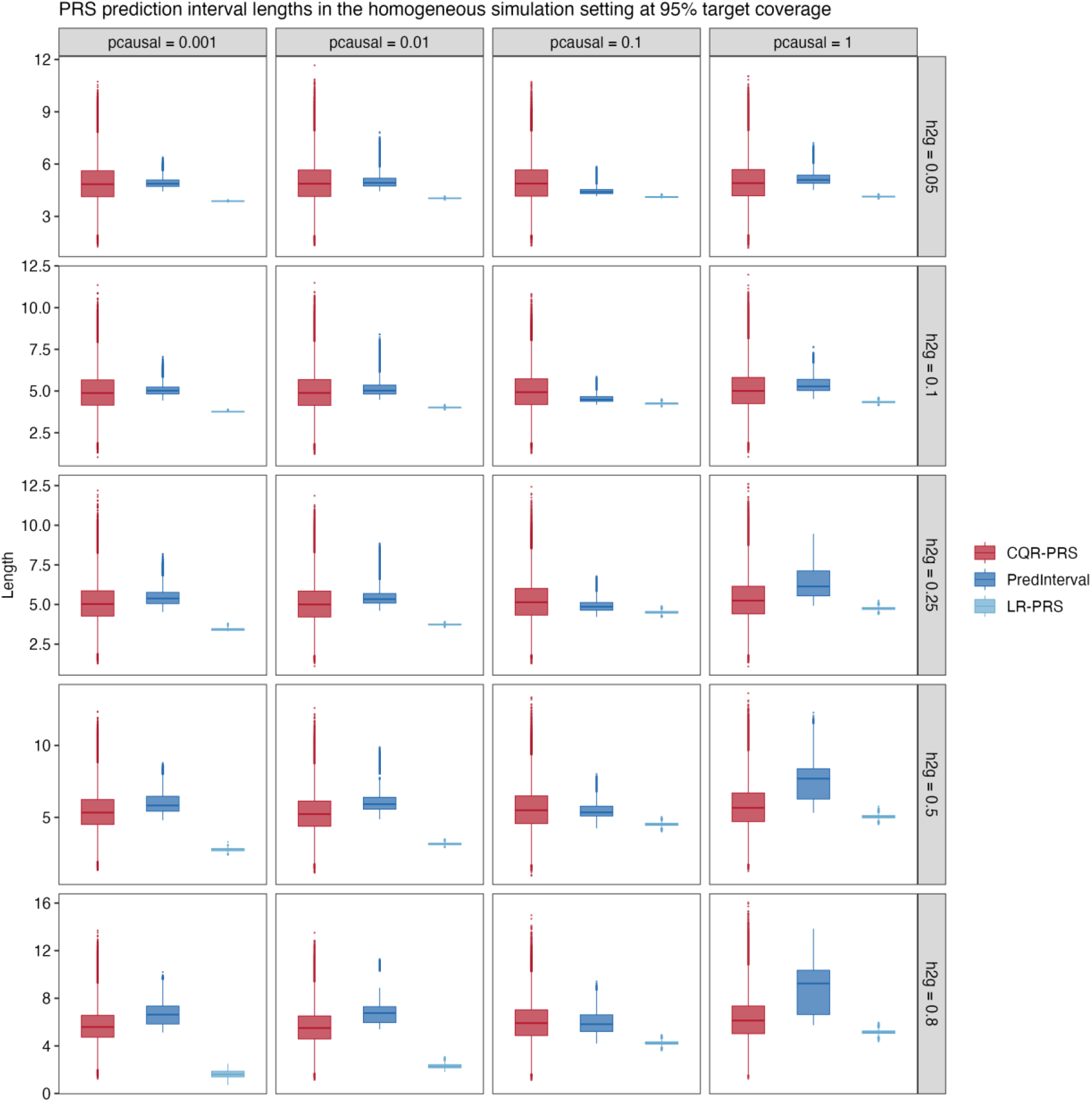
Interval length for CQR-PRS, PredInterval, and LR-PRS intervals in the homogeneous simulation setting, with varying heritability and polygenicity parameters. Each boxplot represents the distribution across a total of 26,377 (individuals in test data) × 20.

**Figure S3:**
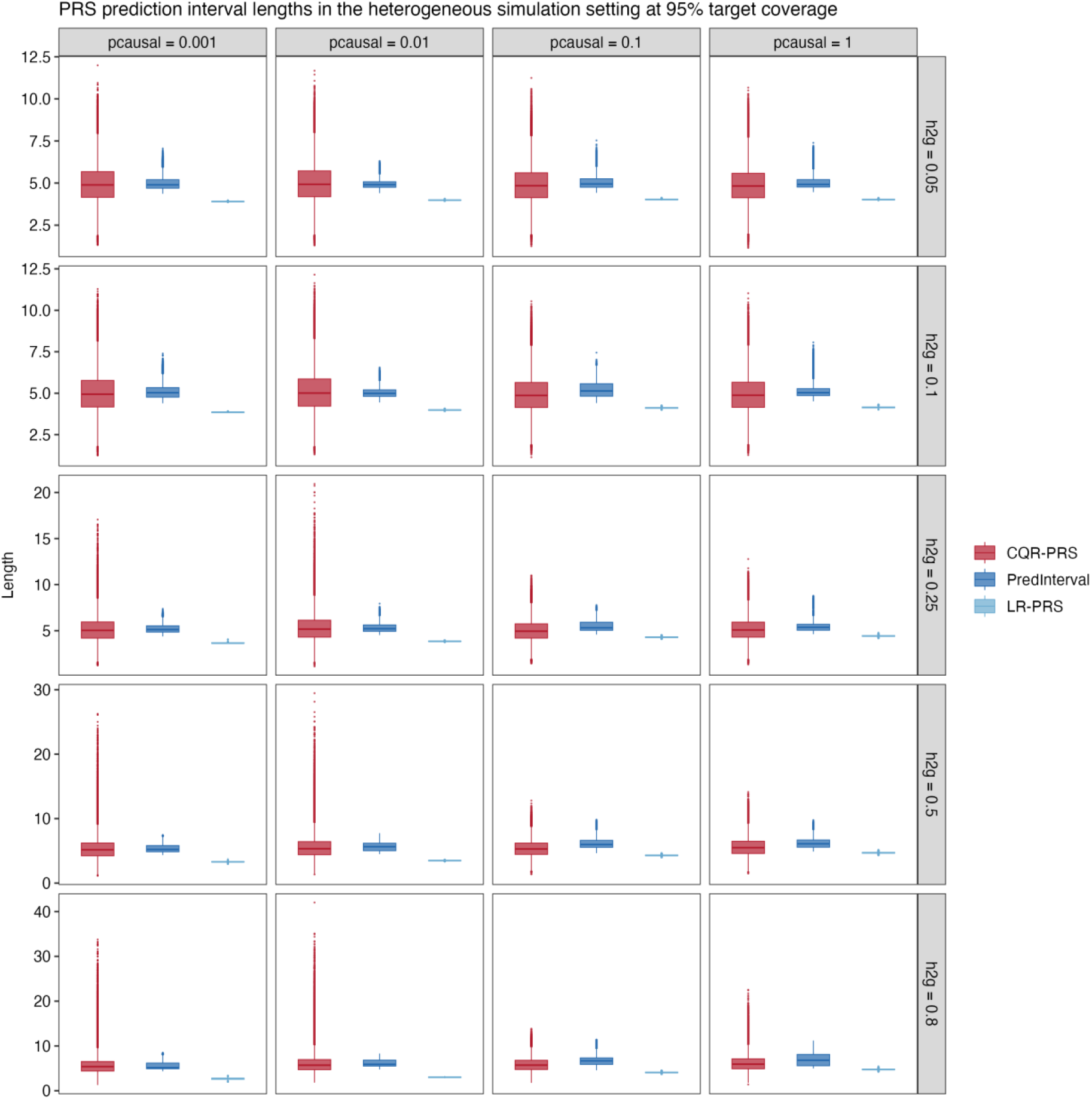
Interval length for CQR-PRS, PredInterval, and LR-PRS intervals in the heterogeneous simulation setting, with varying heritability and polygenicity parameters. Each boxplot represents the distribution across a total of 26,377 (individuals in test data) × 20.

**Figure S4:**
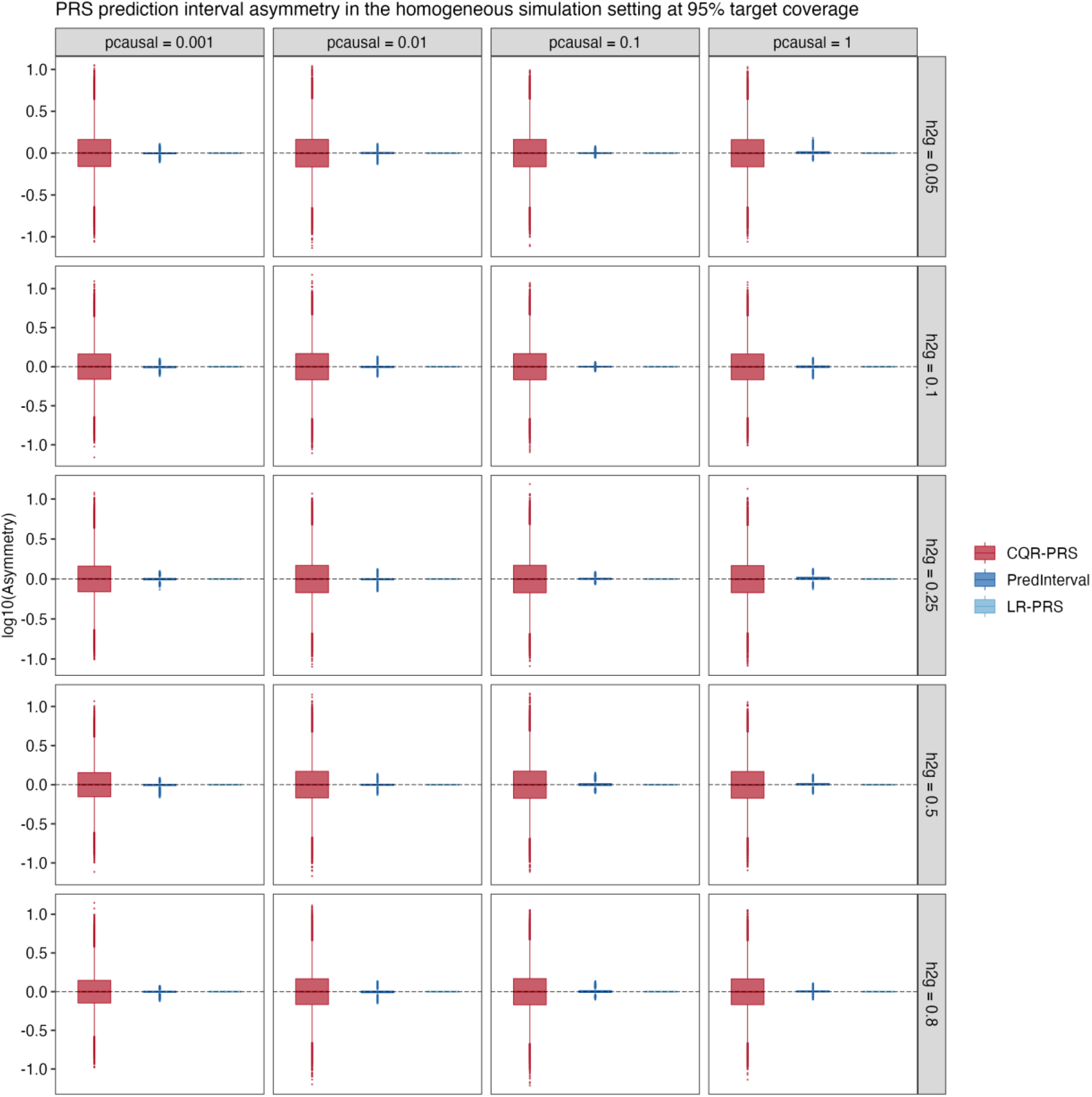
Interval asymmetry for CQR-PRS, PredInterval, and LR-PRS intervals in the homogeneous simulation setting, with varying heritability and polygenicity parameters. Each boxplot represents the distribution across a total of 26,377 (individuals in test data) × 20.

**Figure S5:**
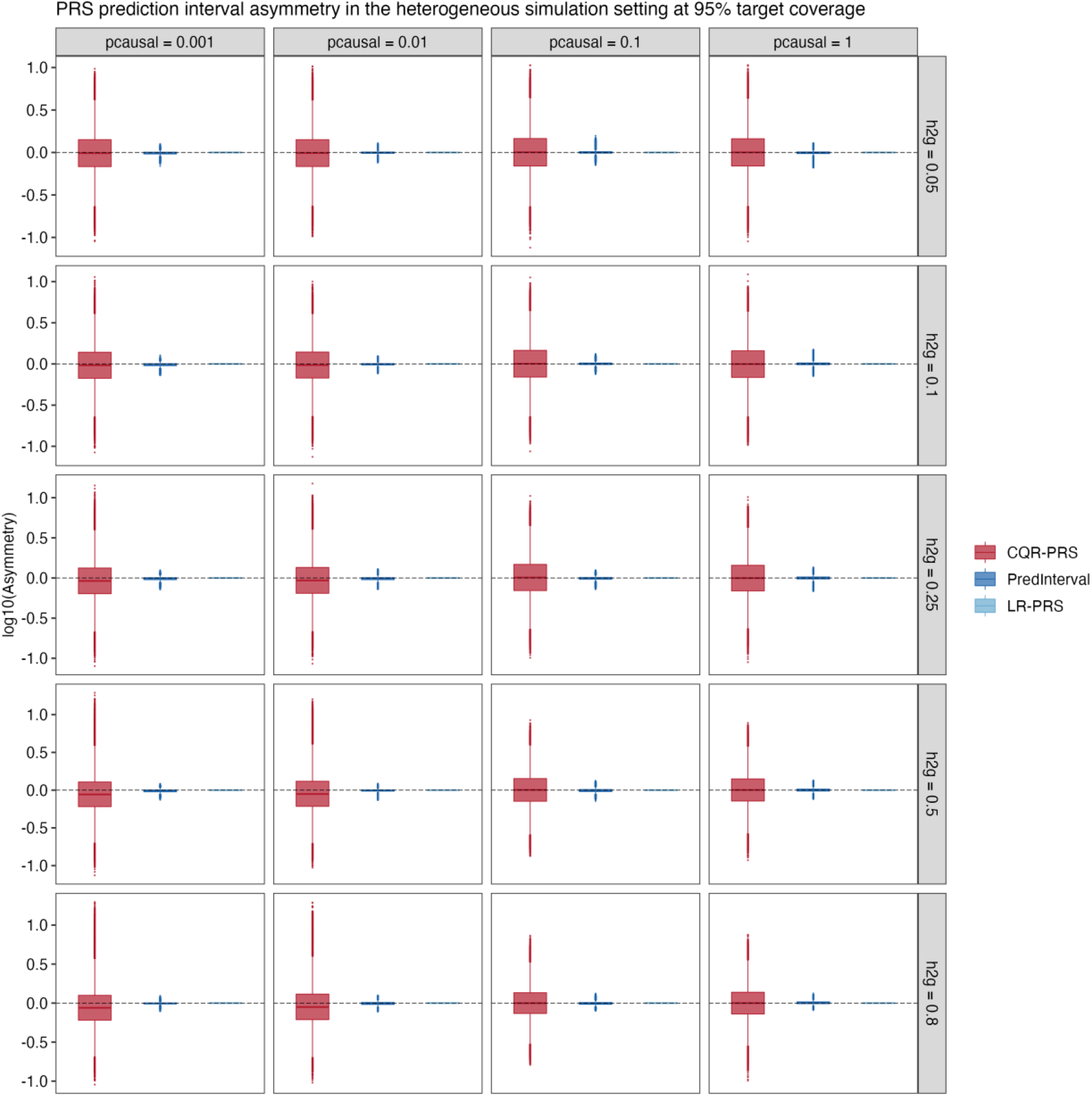
Interval asymmetry for CQR-PRS, PredInterval, and LR-PRS intervals in the heterogeneous simulation setting, with varying heritability and polygenicity parameters. Each boxplot represents the distribution across a total of 26,377 (individuals in test data) × 20.

**Figure S6:**
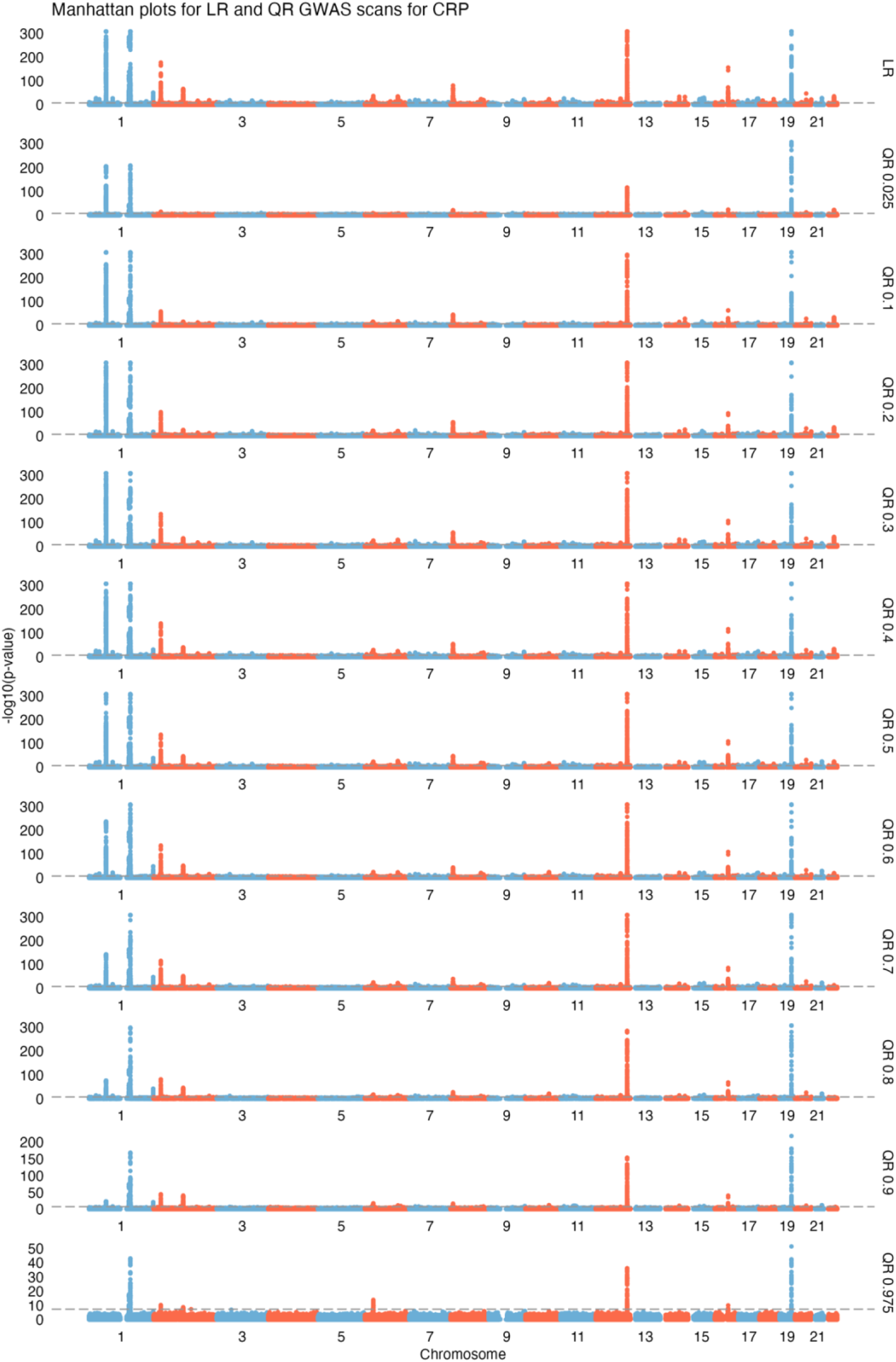
Manhattan plots for LR and QR GWAS (CRP) in UKBB. Quantile regression is run across a grid of quantile levels *τ* ∈ (0.025,0.975).

**Figure S7:**
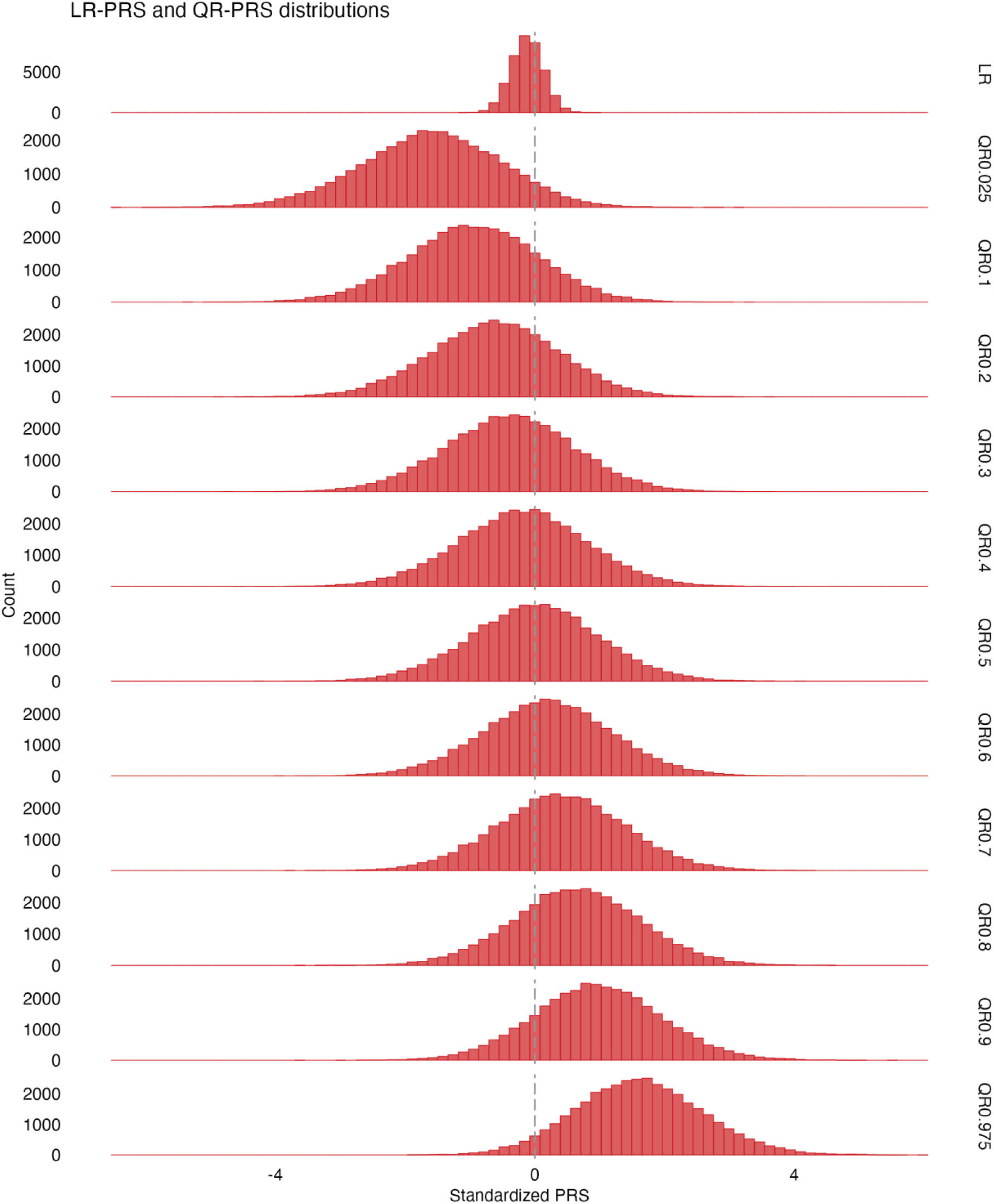
LR-PRS and QR-PRS distribution (CRP) in UKBB. PRS distribution for the mean-based LR-PRS, and individual quantile QR-PRS across a grid of quantile levels *τ* ∈ (0.025, 0.975). The different QR-PRS are standardized with respect to the QR-PRS_0.5_distribution, by subtracting the mean of QR-PRS_0.5_and dividing by the standard deviation.

**Figure S8:**
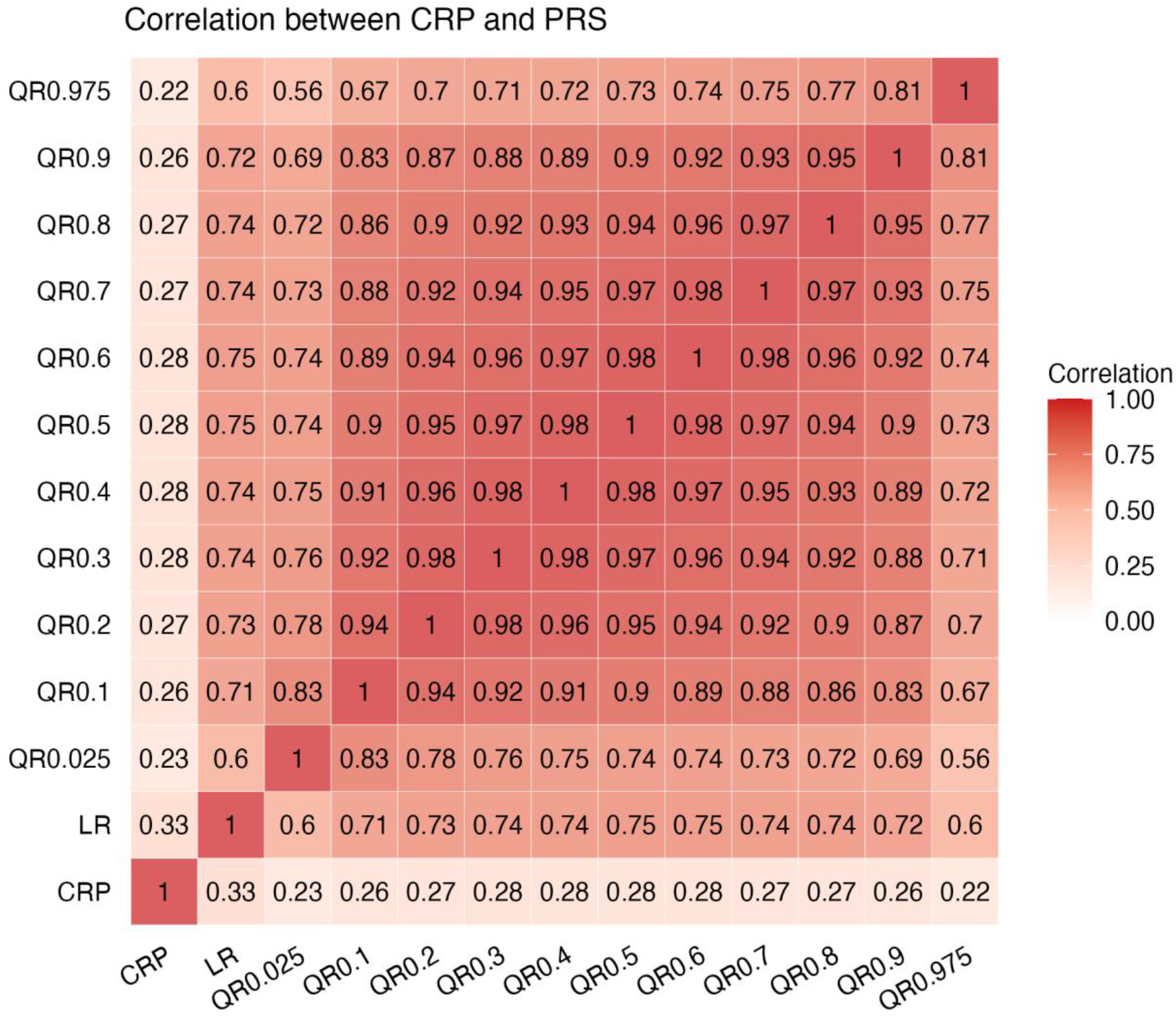
Pearson correlations among actual CRP values, LR-PRS, and QR-PRS across a grid of quantile levels *τ* ∈ (0. 025, 0. 975) in UKBB.

**Figure S9:**
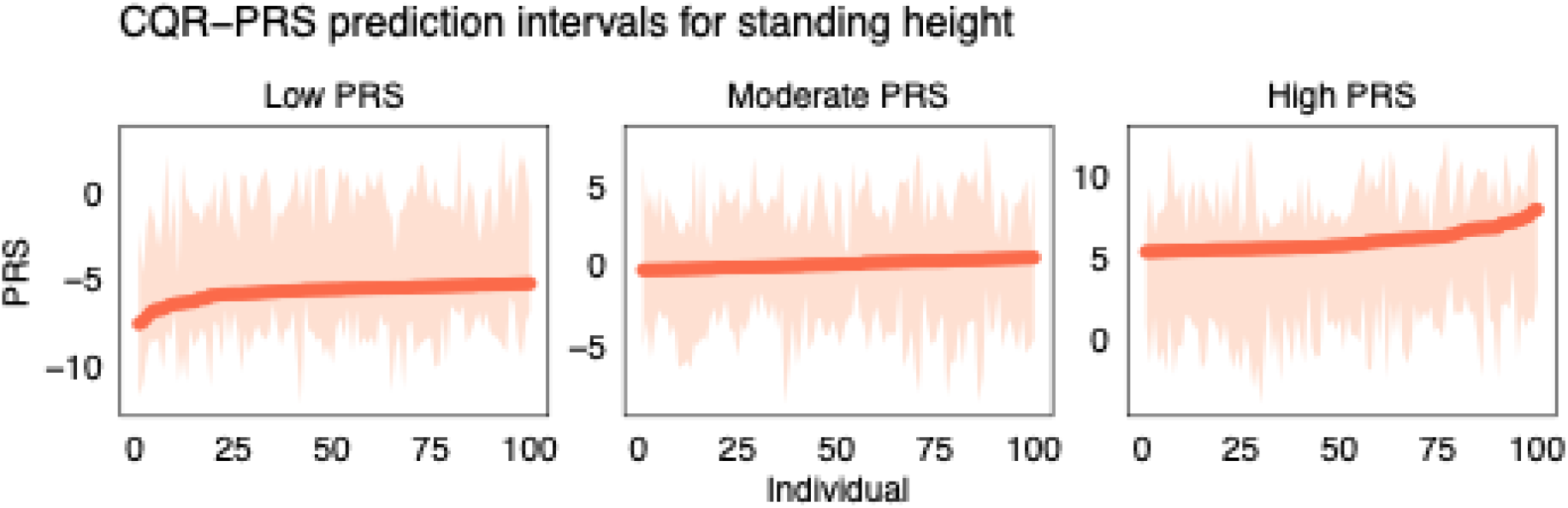
CQR-PRS prediction intervals for standing height. Prediction intervals from CQR-PRS for individuals with low, moderate, and high predicted values. For each group, 100 individuals are selected. Dots correspond to the median QR-PRS_0.5_.

**Figure S10:**
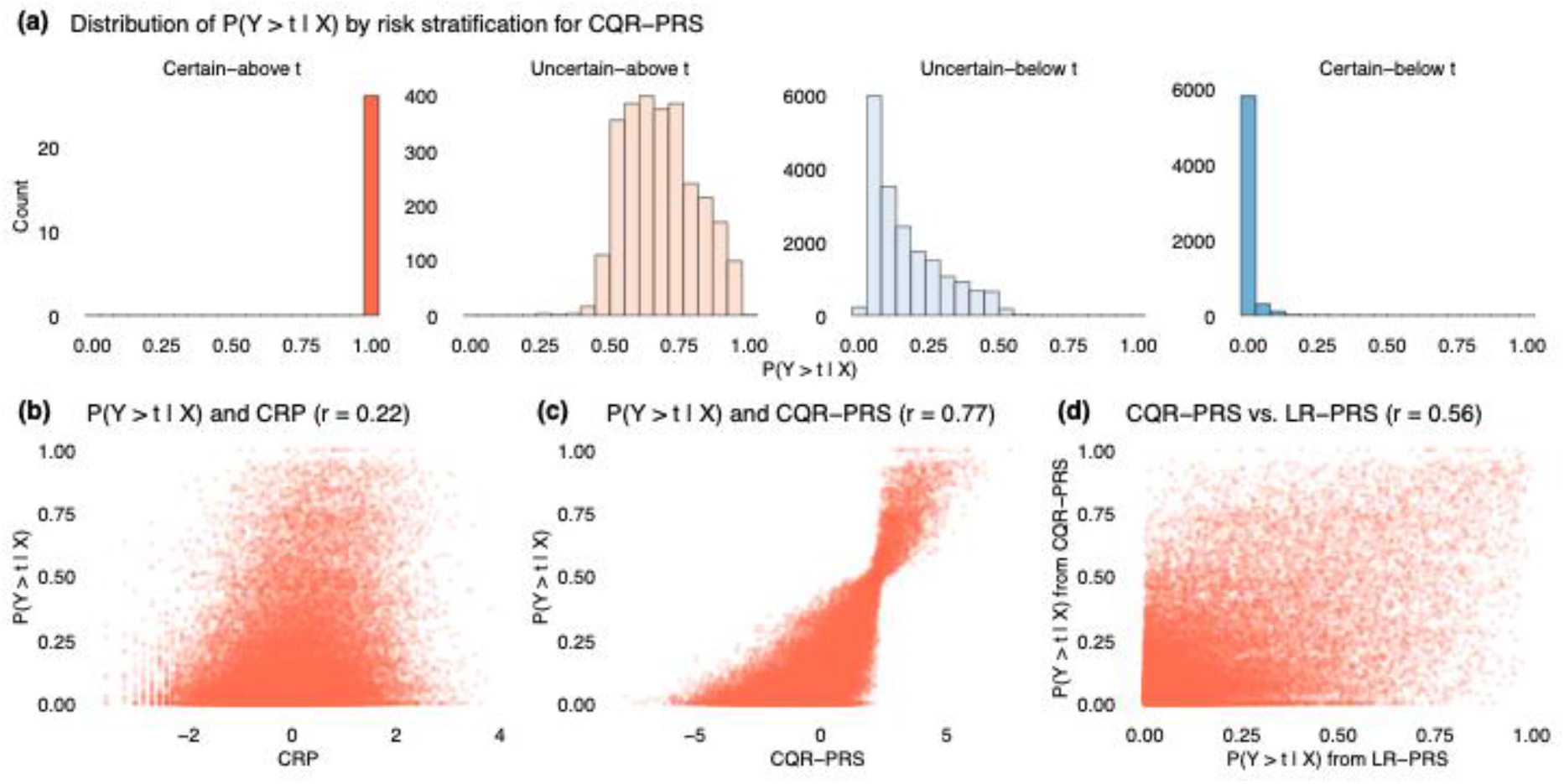
Distribution of *P*(*Y* > *t*|*X*) for individuals in different risk categories and correlation to CRP and QR-PRS_0.5_. Panel **(a)** corresponds to QR-PRS prediction intervals when using 10% as threshold for “high” QR-PRS_0.5_. Panels **(b)**-**(d)** show the relationship between *P*(*Y* > *t*|*X*), CRP, and QR-PRS_0.5_: **(b)** *P*(*Y* > *t*|*X*) and actual CRP, *r* = 0.23; **(c)** *P*(*Y* > *t*|*X*) and QR-PRS_0.5_, *r* = 0.79; **(d)** *P*(*Y* > *t*|*X*) from QR-PRS vs. LR-PRS, *r* = 0.67.

**Figure S11:**
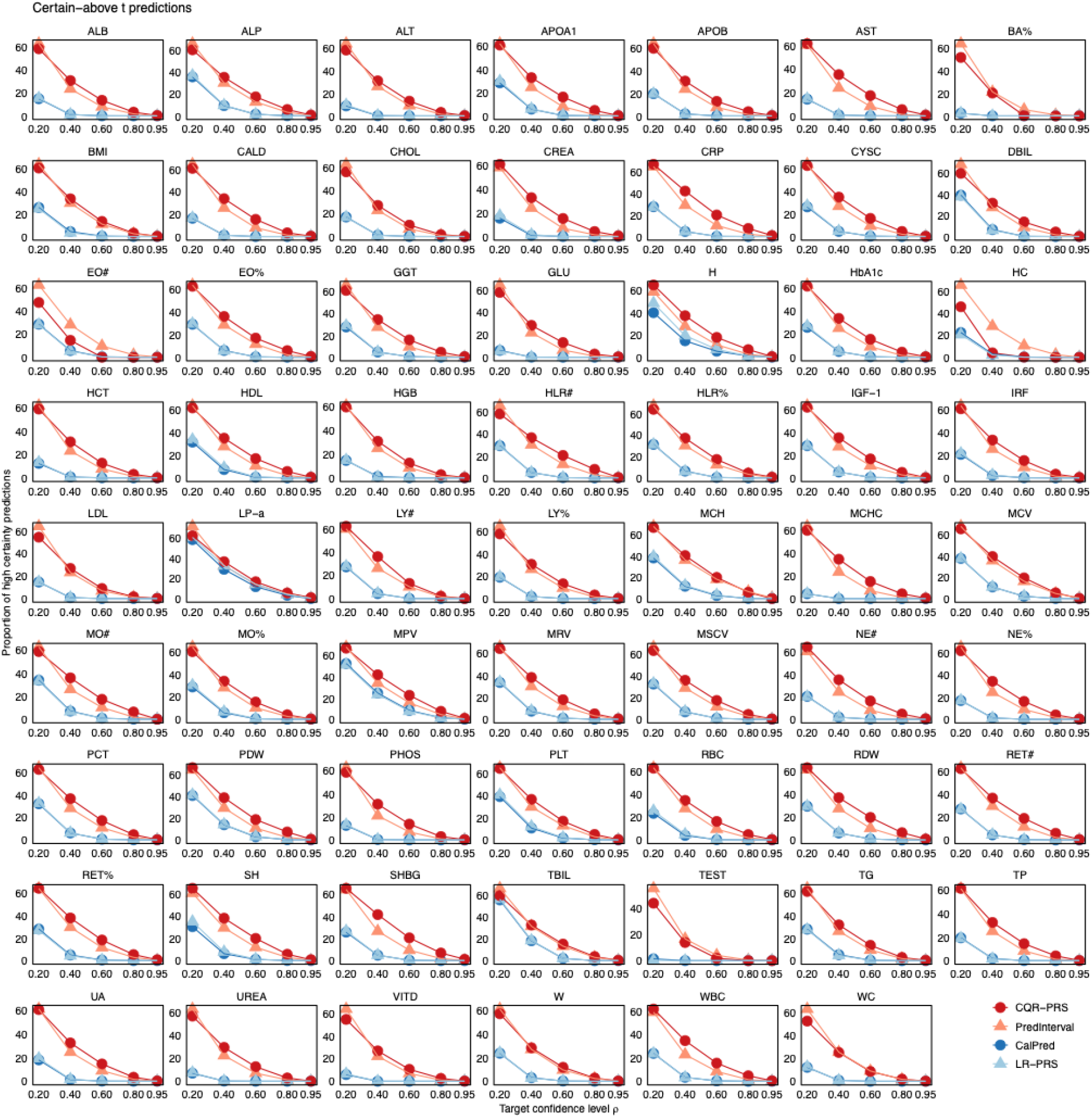
Proportion of certain-above *t* predictions for 62 traits at different target coverage levels (ρ = 0.2, 0.4, 0.6, 0.8, 0.95) in UKBB. Proportion of high certainty predictions is calculated as # certain-above t / # above t. The threshold *t* corresponds to the 90^th^ QR-PRS_0.5_/LR-based PRS percentile.

**Figure S12:**
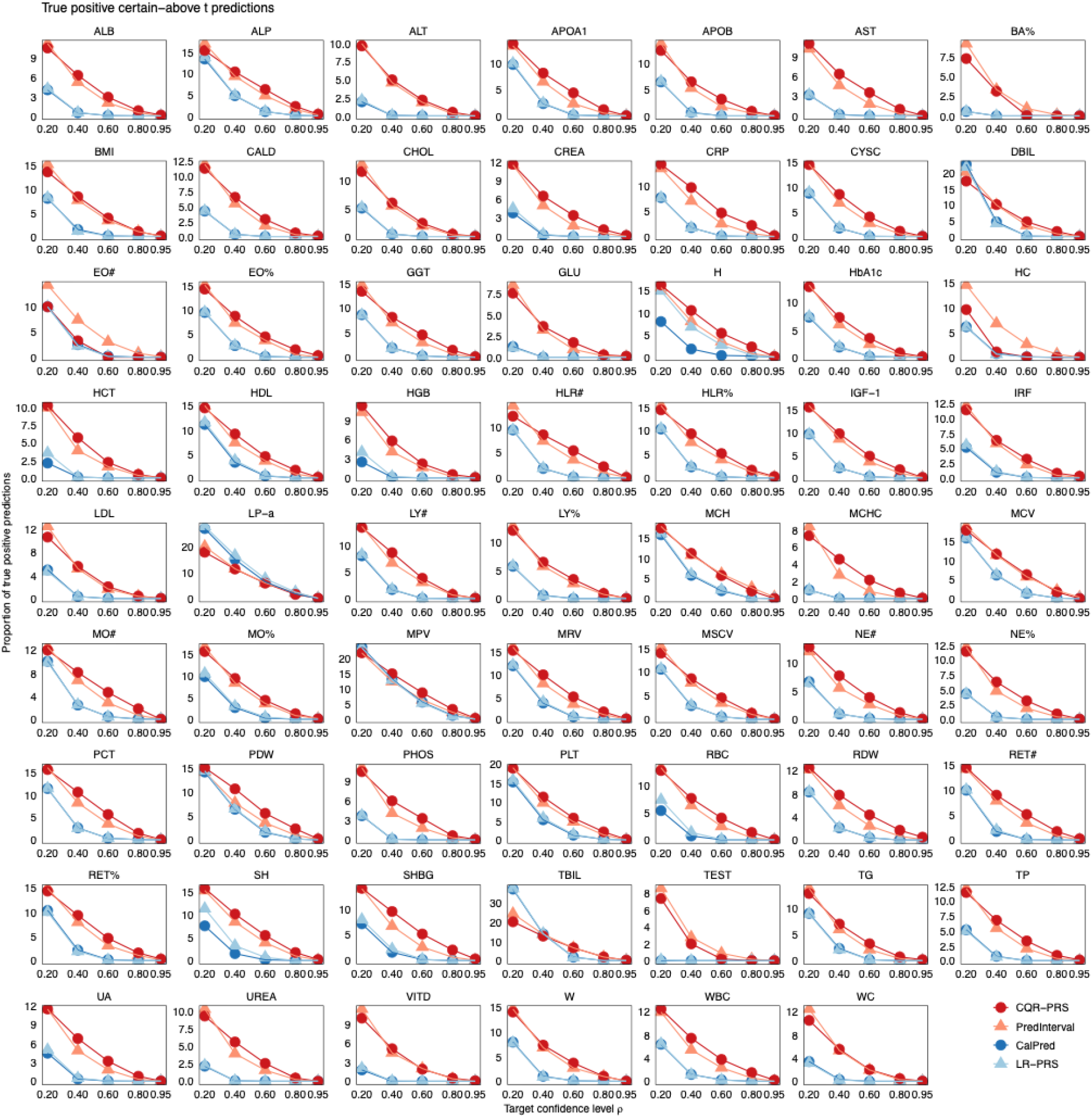
Proportion of true positive certain-above t predictions for 62 traits at different target coverage levels (ρ = 0.2, 0.4, 0.6, 0.8, 0.95) in UKBB. Proportion of true positive predictions is calculated as # (certain-above t & trait above t) / # above t. The threshold *t* corresponds to the 90^th^ QR-PRS_0.5_/LR-based PRS or trait percentile depending on context.

**Figure S13:**
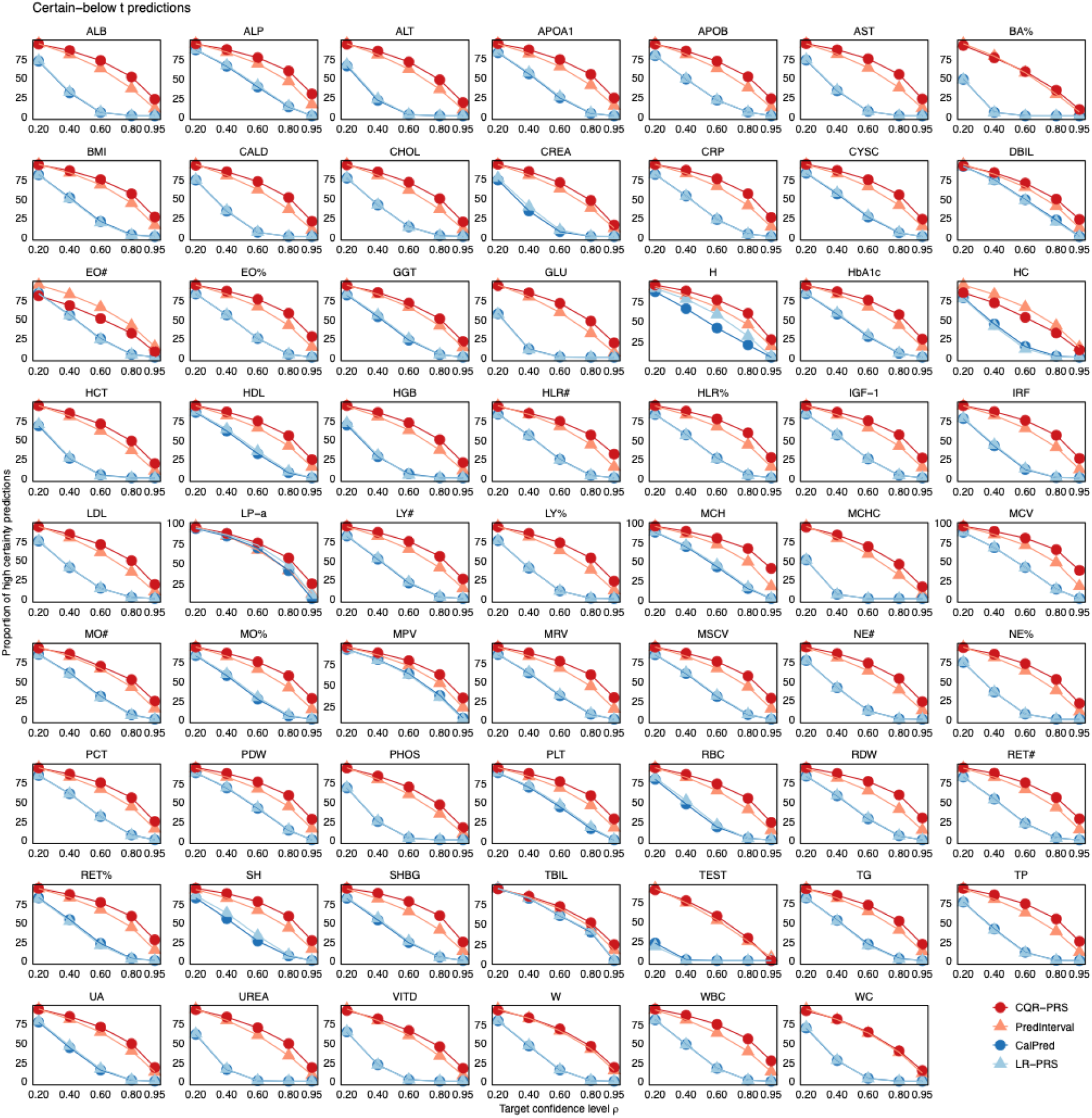
Proportion of certain-below t predictions for 62 traits at different target coverage levels (ρ = 0.2, 0.4, 0.6, 0.8, 0.95) in UKBB. Proportion of high certainty predictions is calculated as # certain-below t / # below t. The threshold *t* corresponds to the 90^th^ QR-PRS_0.5_/LR-based PRS percentile.

**Figure S14:**
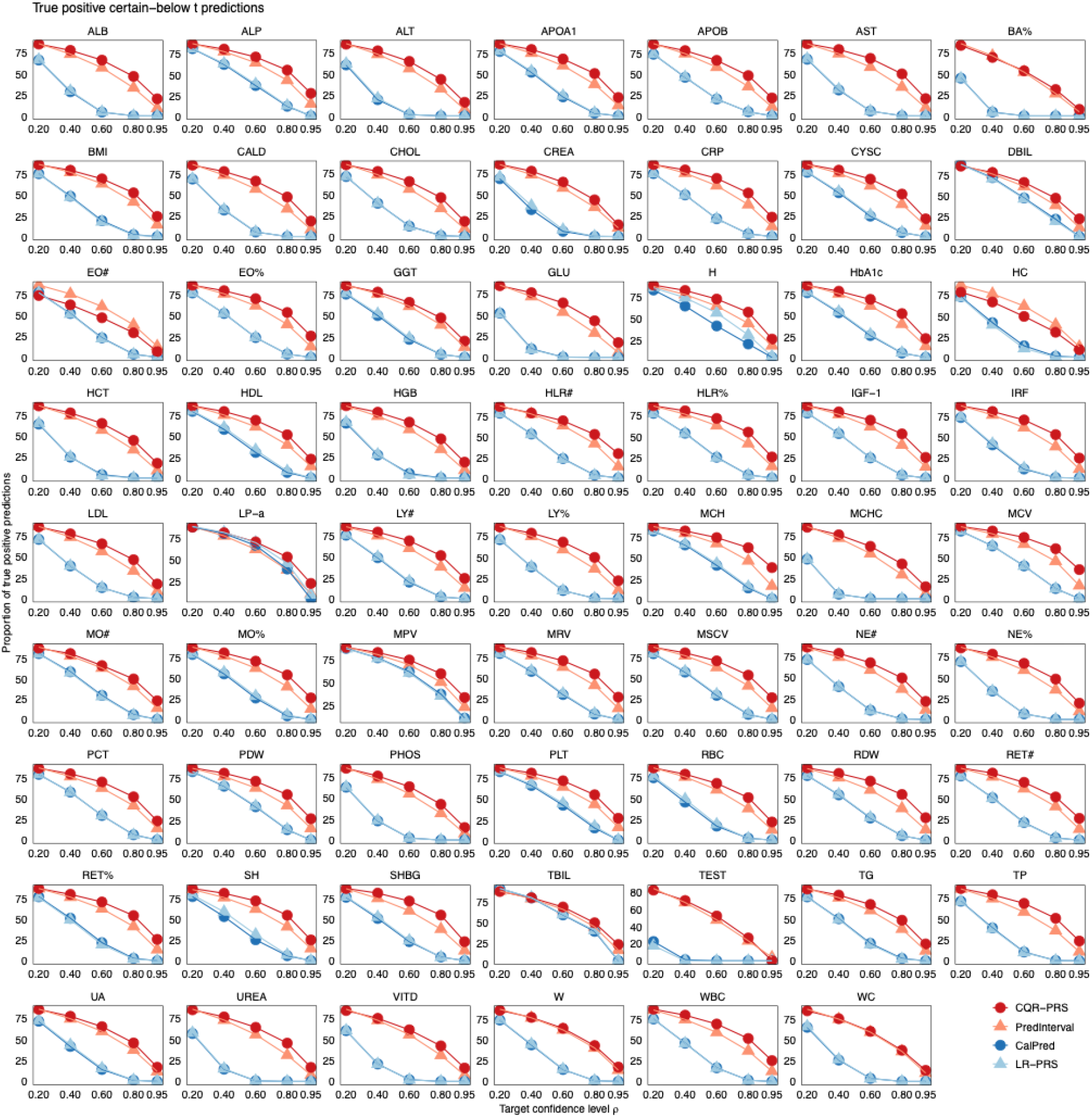
Proportion of true positive certain-below t predictions for 62 traits at different target coverage levels (ρ = 0.2, 0.4, 0.6, 0.8, 0.95) in UKBB. Proportion of true positive predictions is calculated as # (certain-below t & trait below t) / # below t. The threshold *t* corresponds to the 90^th^ QR-PRS_0.5_/LR-based PRS or trait percentile depending on context.

**Figure S15:**
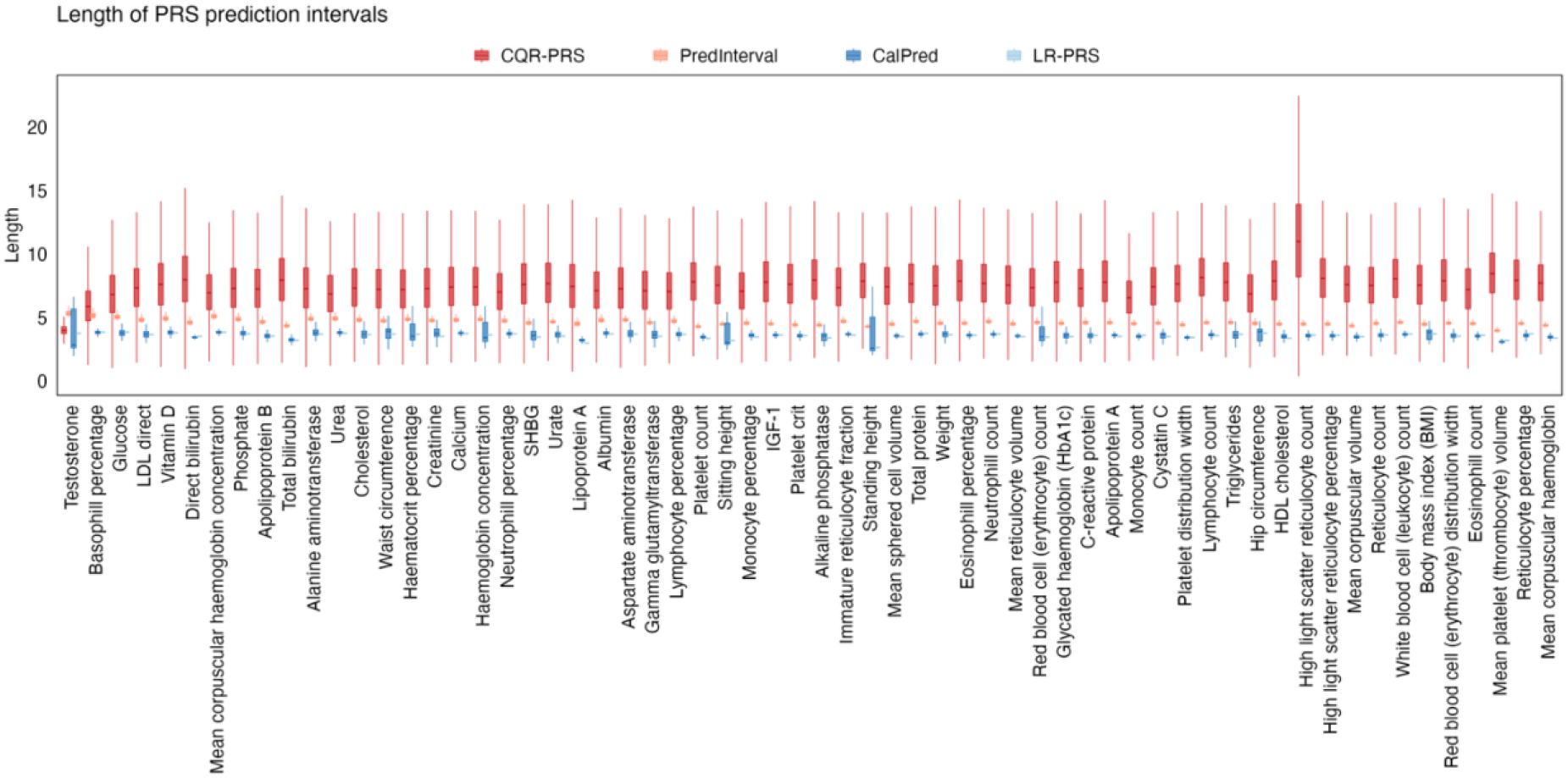
Interval length for CQR-PRS, PredInterval, CalPred and LR-PRS intervals across 62 traits in UKBB.

**Figure S16:**
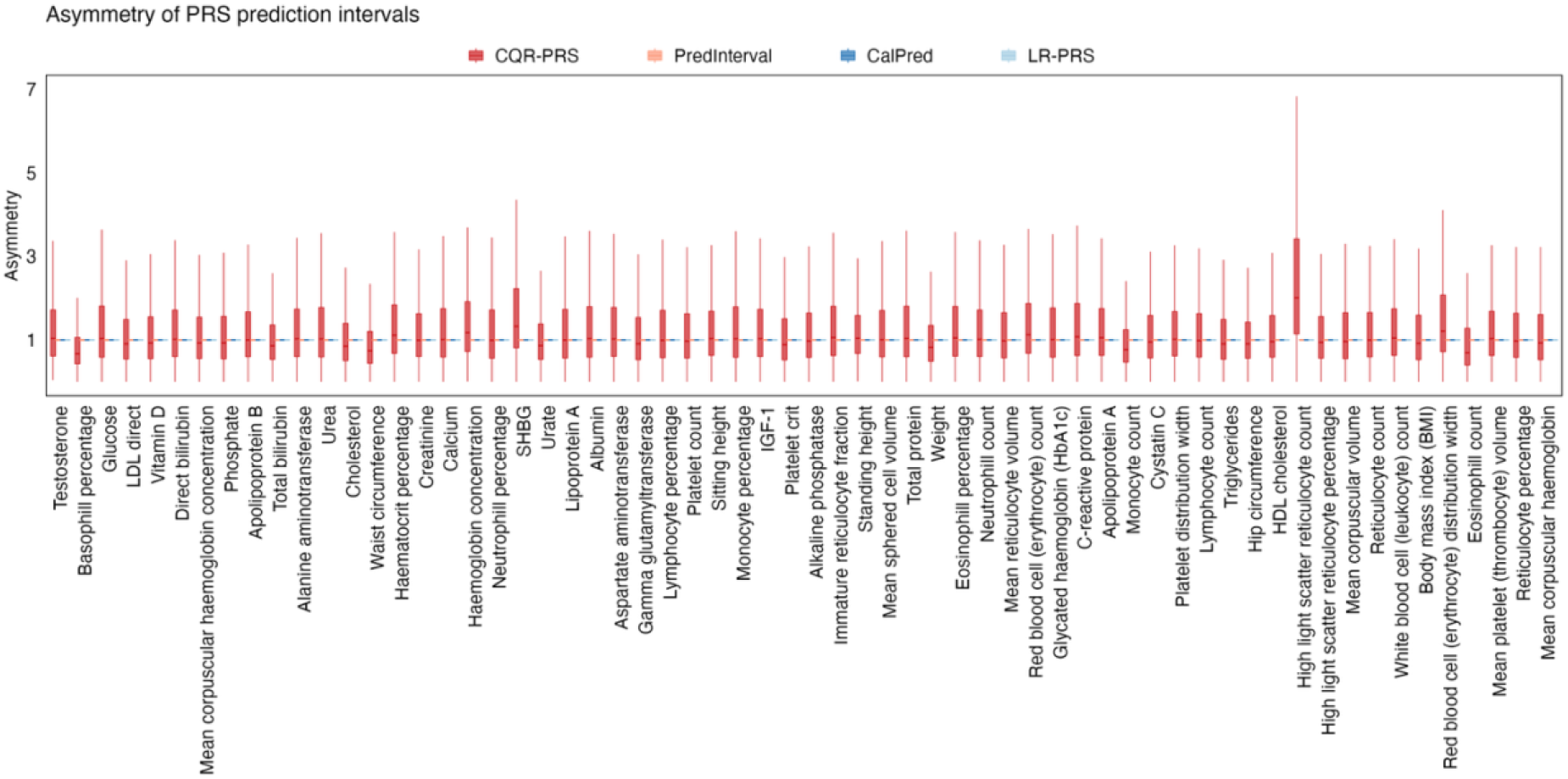
Interval asymmetry for CQR-PRS, PredInterval, CalPred and LR-PRS intervals across 62 traits in UKBB.

**Figure S17:**
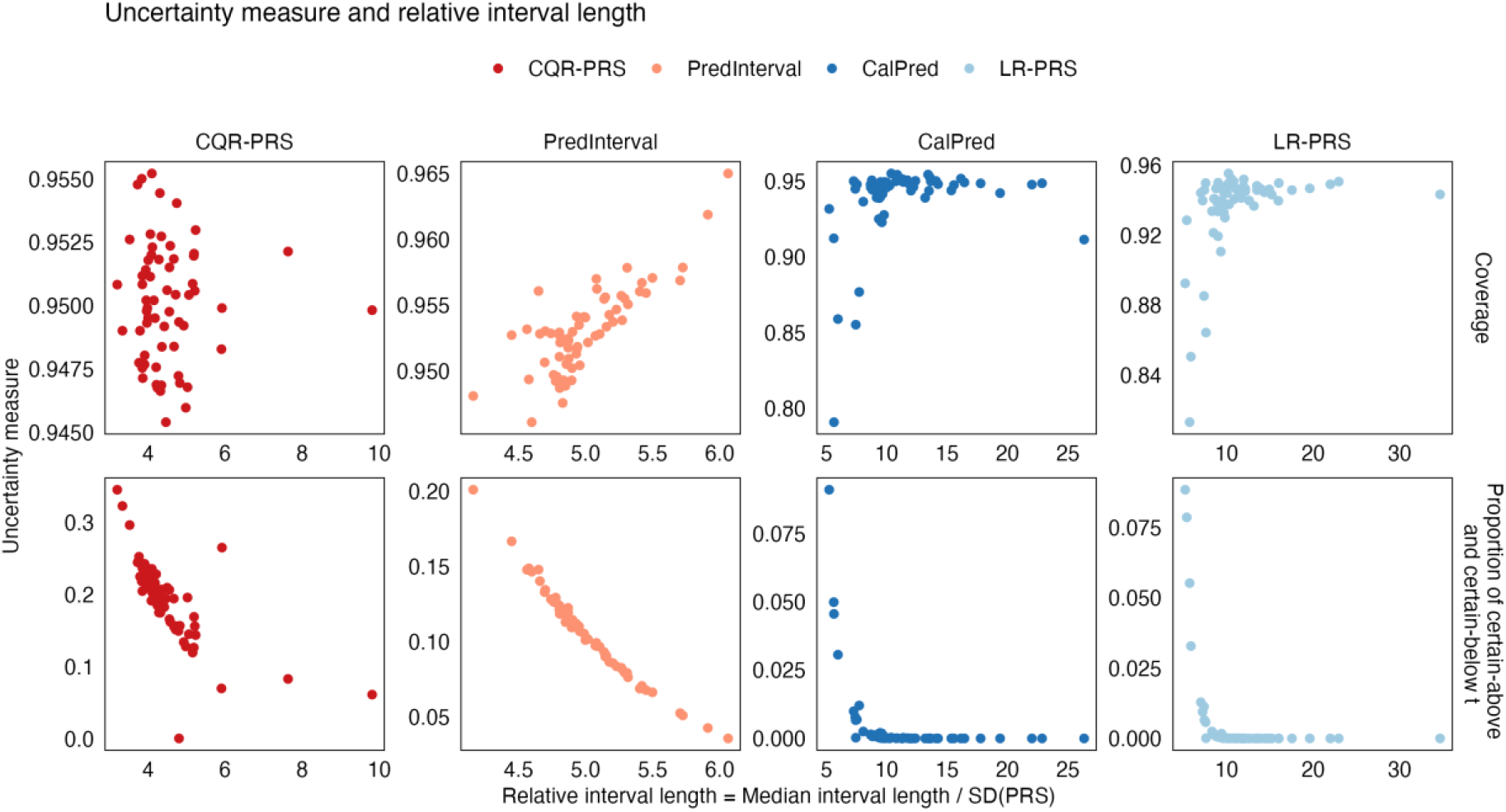
Performance of CQR-PRS, PredInterval, CalPred and LR-PRS evaluated by uncertainty measures and relative interval length. The top row illustrates empirical coverage, while the bottom row displays the proportion of certain predictions (representing individuals whose prediction intervals fall strictly above or below a defined threshold *t*). Both metrics are plotted against the relative interval length on the x-axis, which is standardized as the median interval length divided by the standard deviation of the PRS.

**Figure S18:**
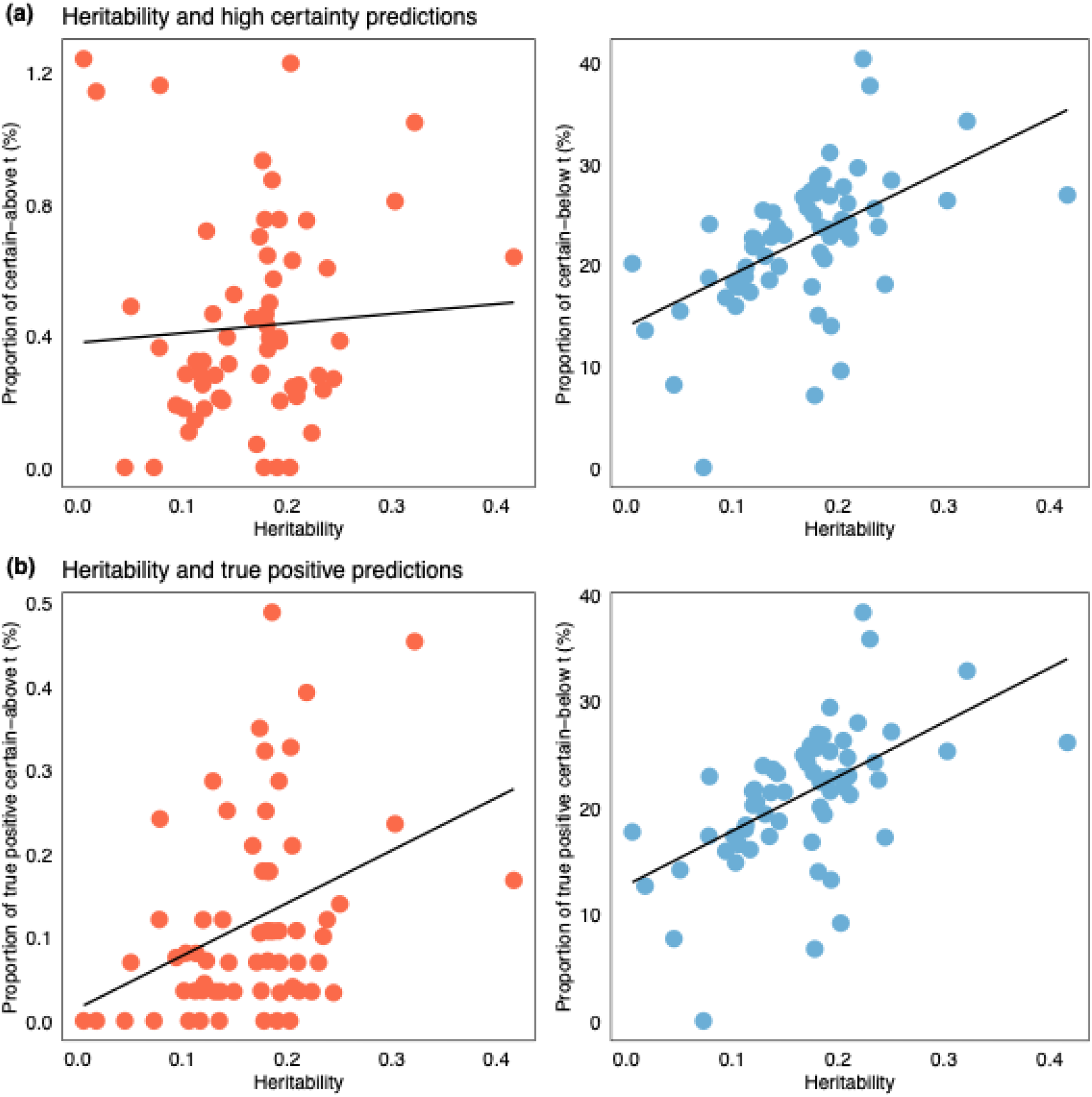
Relationship between heritability for a trait and uncertainty quantification in UKBB. **(a)** Heritability and proportion of high certainty predictions. Left: certain-above t (*r* = 0.06, *p* = 0.61). Right: certain-below t (*r* = 0.53, *p* = 8.4E-6). Proportion of high certainty predictions is calculated as # certain-above t / # above t and # certain-below t / # below t. **(b)** Heritability and proportion of true positive predictions. Left: certain-above t (*r* = 0.37, *p* = 2.7E-3). Right: certain-below t (*r* = 0.55, *p* = 3.2E-6). Proportion of true positive predictions is calculated as # (certain-above t & trait above t) / # above t and # (certain-below t & trait below t) / # below t. SNP heritability was estimated by LDSC. Prediction uncertainty was quantified using CQR-PRS prediction intervals. The threshold *t* corresponds to the 90^th^ percentile of QR-PRS_0.5_for each trait.

**Figure S19:**
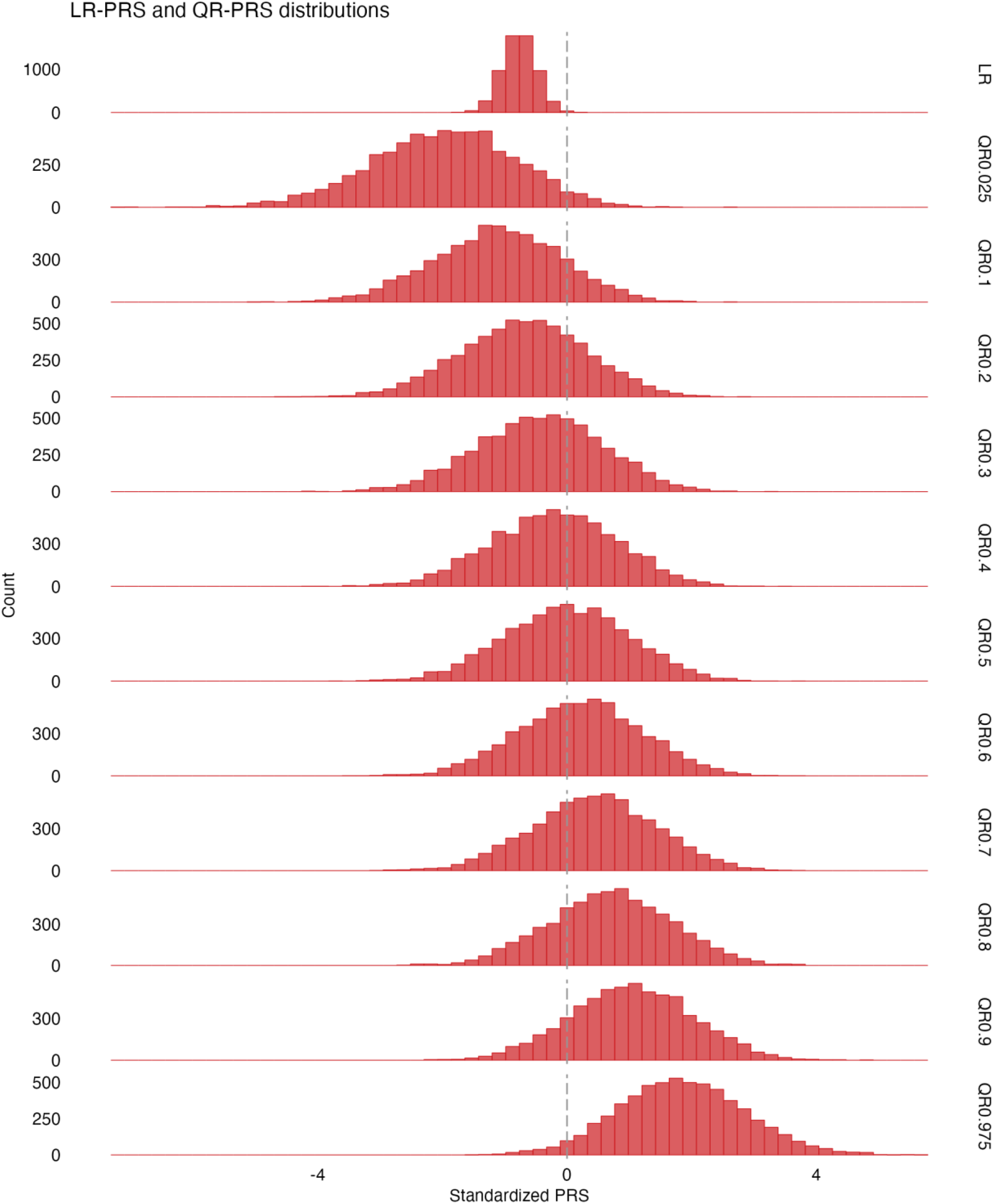
LR-PRS and QR-PRS distribution (CRP) in ProgeNIA/SardiNIA. PRS distribution for the mean-based LR-PRS, and individual quantile QR-PRS across a grid of quantile levels *τ* ∈ (0.025, 0.975). The different QR-PRS are standardized with respect to the QR-PRS_0.5_distribution, by subtracting the mean of QR-PRS_0.5_and dividing by the standard deviation.

**Figure S20:**
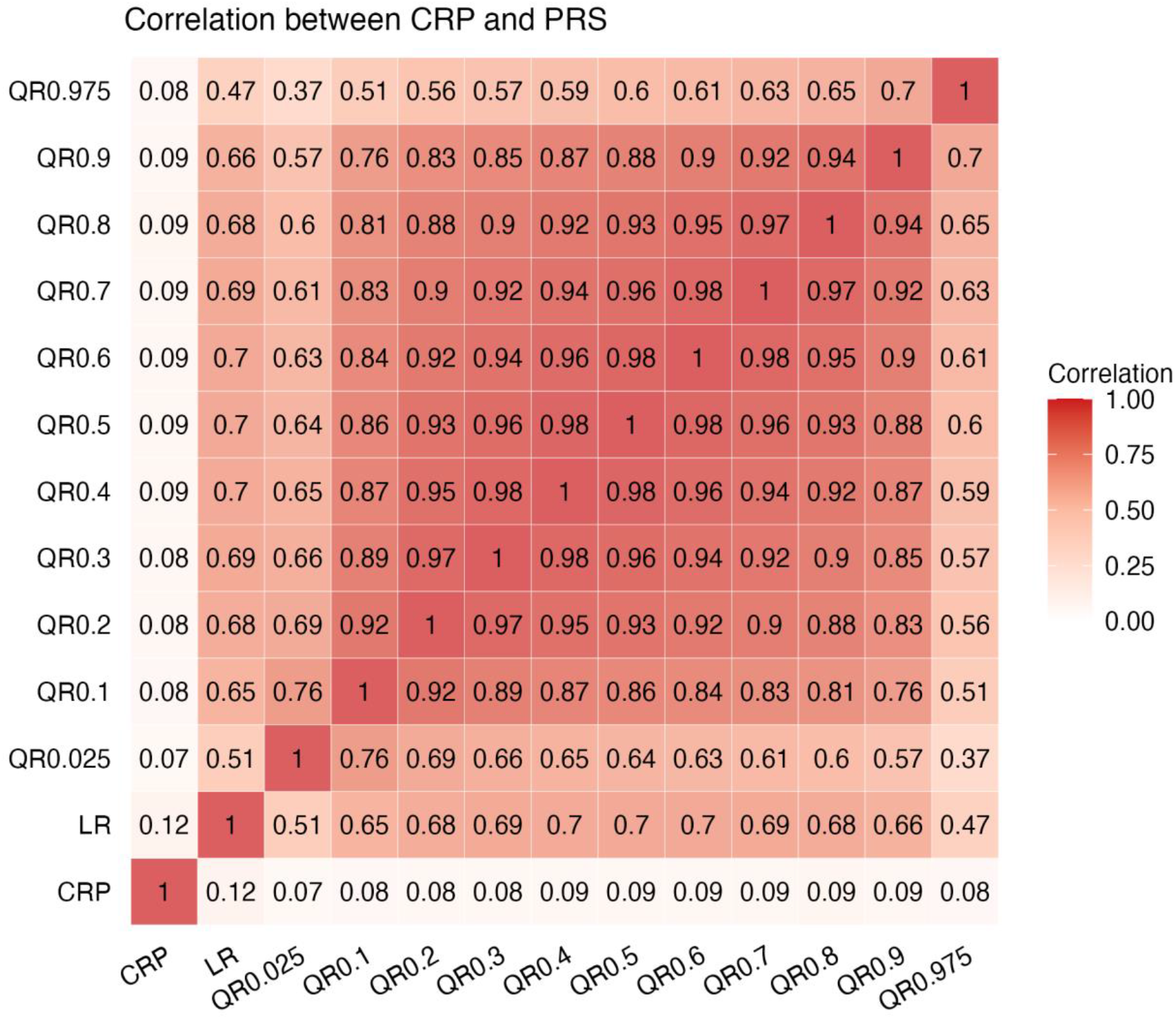
Pearson correlations among actual CRP values, LR-PRS, and QR-PRS across a grid of quantile levels *τ* ∈ (0. 025, 0. 975) in ProgeNIA/SardiNIA.

**Figure S21:**
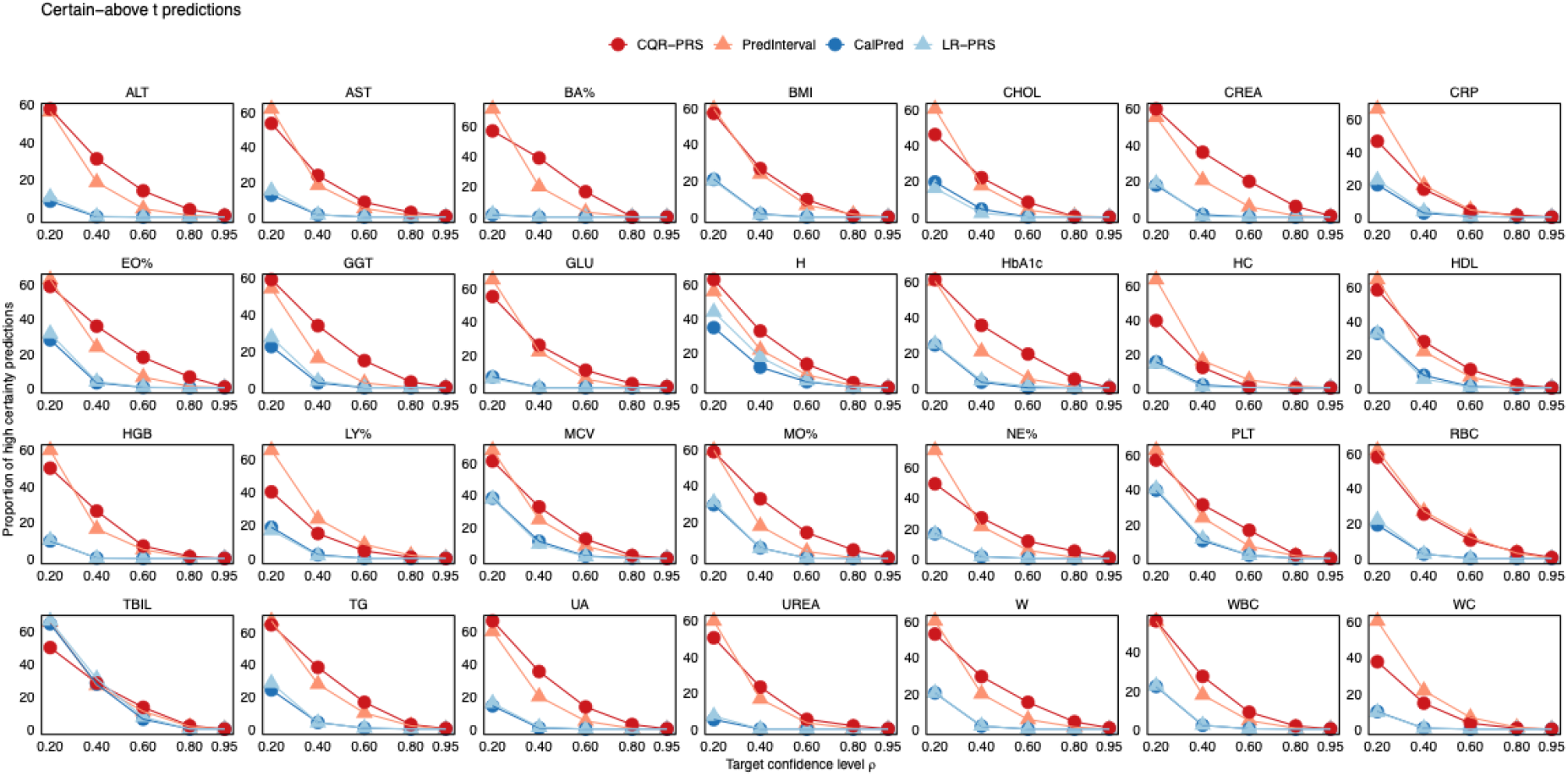
Proportion of certain-above t predictions for 28 traits at different target coverage levels (ρ = 0.2, 0.4, 0.6, 0.8, 0.95) in ProgeNIA/SardiNIA. Proportion of high certainty predictions is calculated as # certain-above t / # above t. The threshold *t* corresponds to the 90^th^ QR-PRS_0.5_/LR-based PRS percentile.

**Figure S22:**
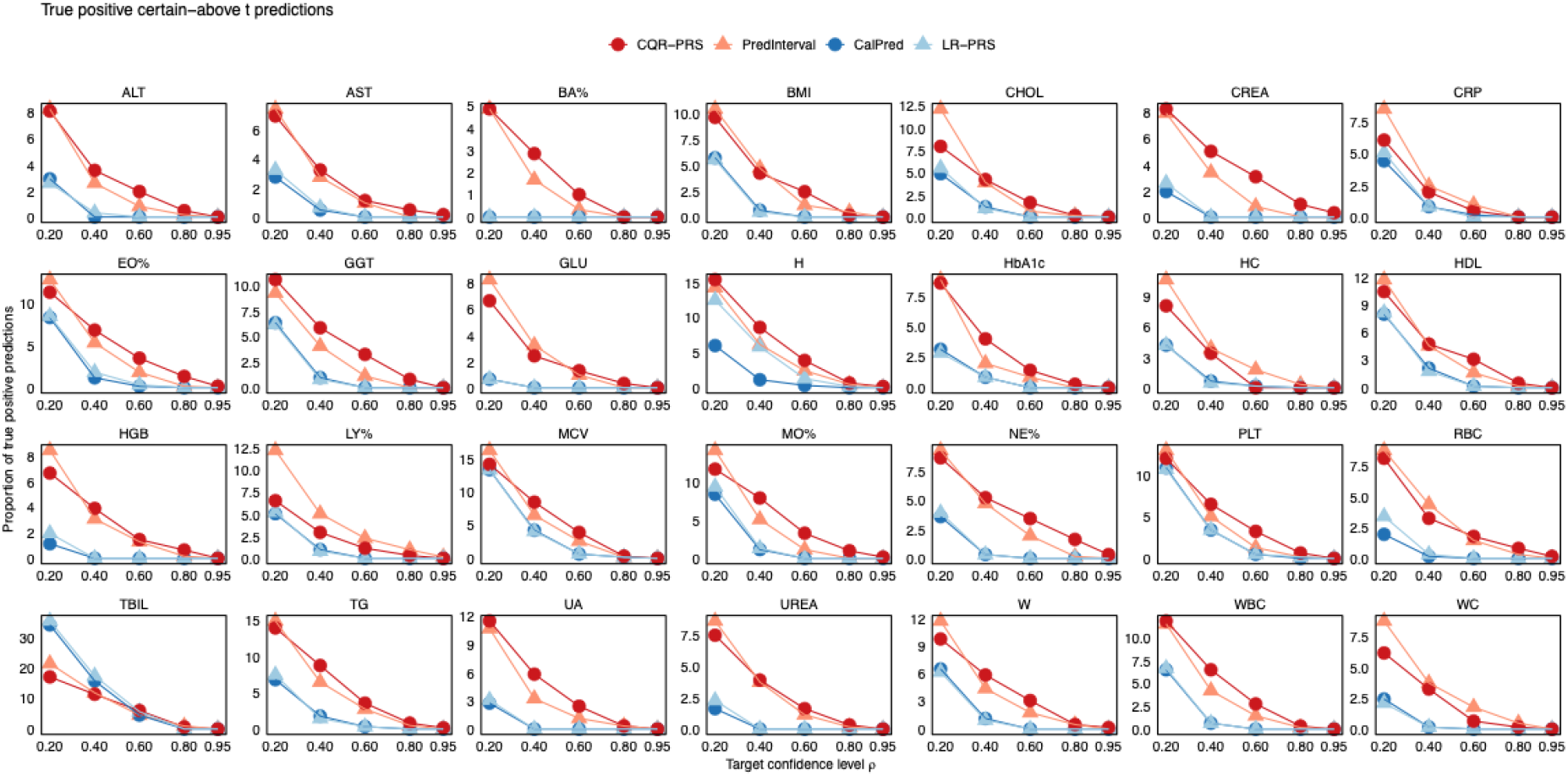
Proportion of true positive certain-above t predictions for 28 traits at different target coverage levels (ρ = 0.2, 0.4, 0.6, 0.8, 0.95) in ProgeNIA/SardiNIA. Proportion of true positive predictions is calculated as # (certain-above t & trait above t) / # above t. The threshold *t* corresponds to the 90^th^ QR-PRS_0.5_/LR-based PRS or trait percentile depending on context.

**Figure S23:**
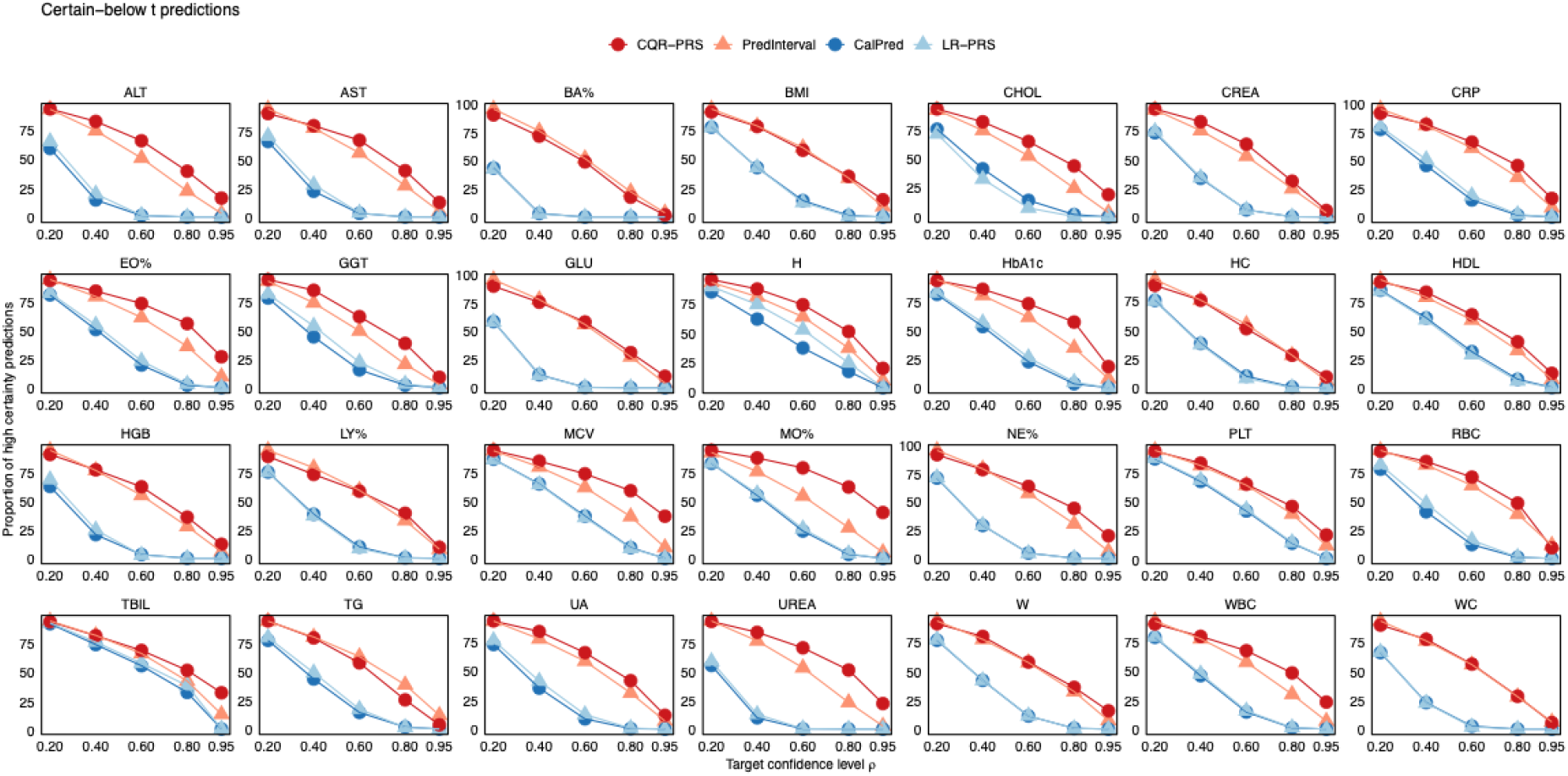
Proportion of certain-below t predictions for 28 traits at different target coverage levels (ρ = 0.2, 0.4, 0.6, 0.8, 0.95) in ProgeNIA/SardiNIA. Proportion of high certainty predictions is calculated as # certain-below t / # below t. The threshold *t* corresponds to the 90^th^ QR-PRS_0.5_/LR-based PRS percentile.

**Figure S24:**
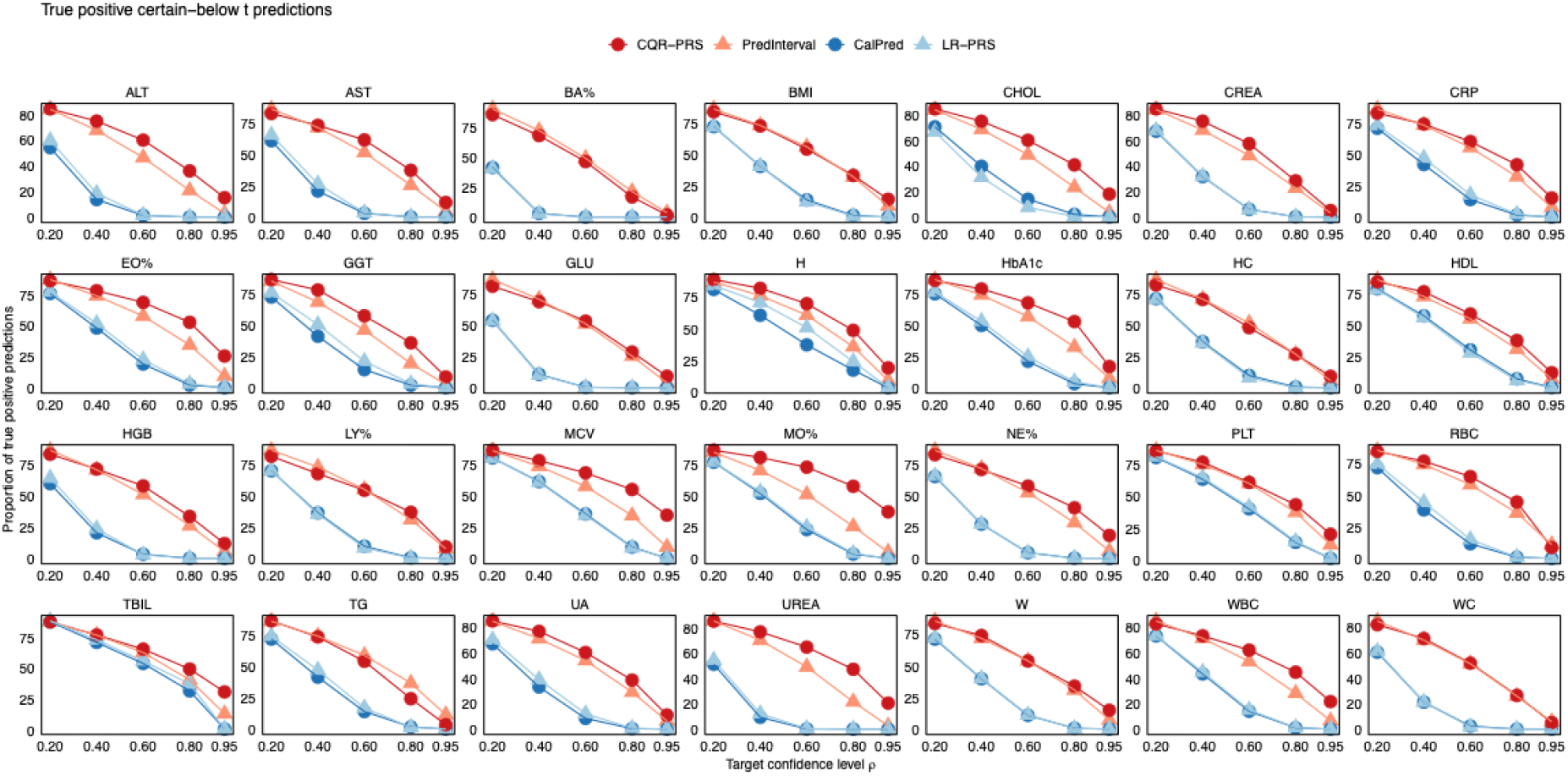
Proportion of true positive certain-below t predictions for 28 traits at different target coverage levels (ρ = 0.2, 0.4, 0.6, 0.8, 0.95) in ProgeNIA/SardiNIA. Proportion of true positive predictions is calculated as # (certain-below t & trait below t) / # below t. The threshold *t* corresponds to the 90^th^ QR-PRS_0.5_/LR-based PRS or trait percentile depending on context.

**Table S1:**
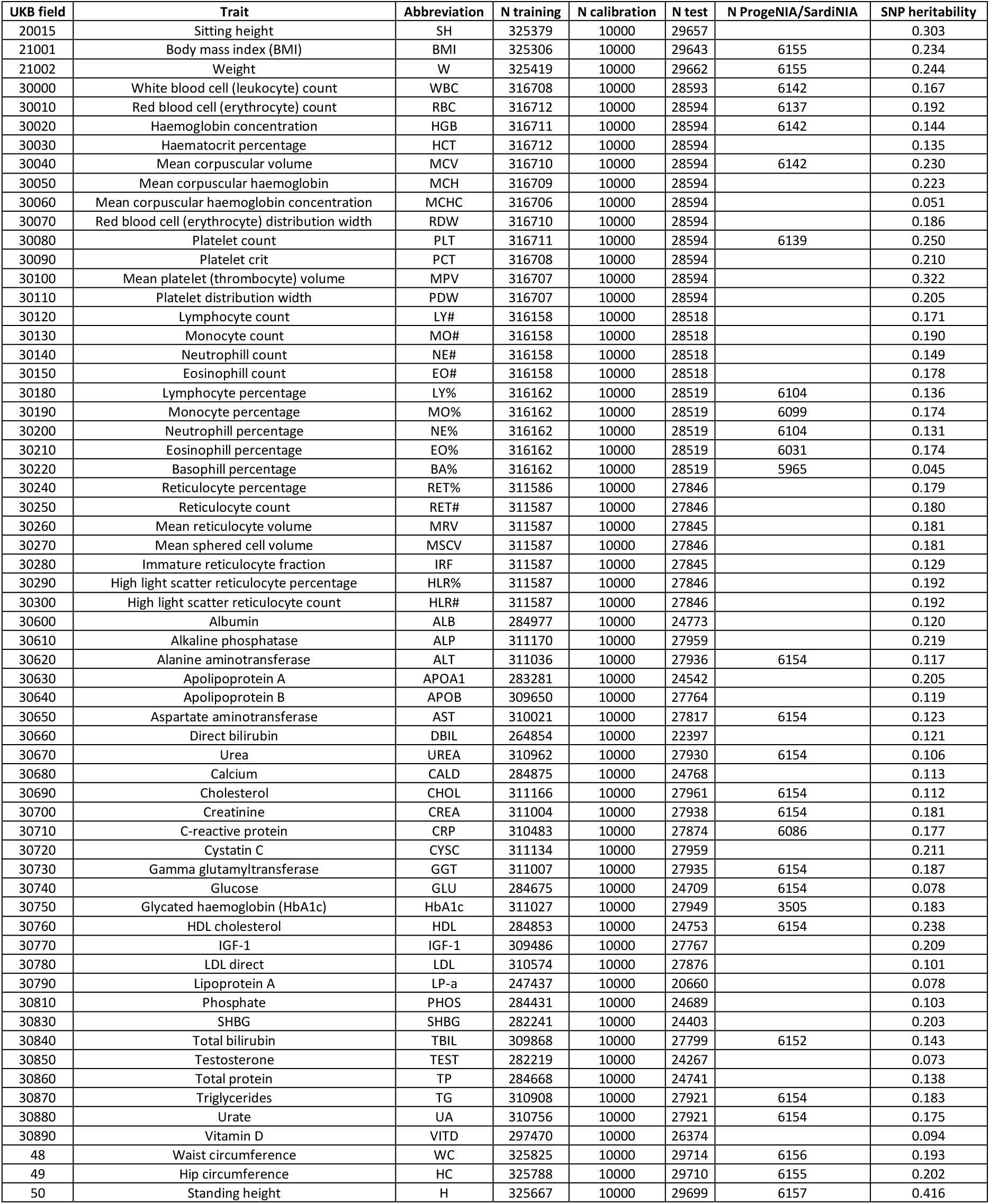
Sample sizes for training, calibration/validation, and test datasets and SNP heritability for each of 62 traits in UKBB, and sample sizes for each of 28 traits in ProgeNIA/SardiNIA. SNP heritability for each trait was estimated using LDSC.

## Notes

### Competing Interest Statement

The authors have declared no competing interest.

### Summary of Updates

This revision includes additional empirical comparisons with recent methods for prediction interval construction.

